# A direct lateral entorhinal cortex to hippocampal CA2 circuit conveys social information required for social memory

**DOI:** 10.1101/2021.04.15.440048

**Authors:** Jeffrey Lopez-Rojas, Christopher A. de Solis, Felix Leroy, Eric R. Kandel, Steven A. Siegelbaum

## Abstract

The storage of information by the hippocampus in long-term memory is thought to involve two distinct but related processes. First, the hippocampus determines whether a given stimulus is novel or familiar; next, the hippocampus stores the novel information in long-term memory. To date, the neural circuits that detect novelty and their relation to the circuits that store information of a specific memory are poorly understood. Here we address this question by examining the circuits by which the CA2 region of the hippocampus, which is essential for social memory, both detects social novelty and stores social memory. CA2, like the more thoroughly studied CA1 region, receives its major excitatory input from the entorhinal cortex through both a direct monosynaptic and indirect trisynaptic pathway. We find that the direct inputs to CA2 from the lateral entorhinal cortex, but not the indirect trisynaptic inputs, provide social information that is required for social memory. However, these direct inputs fail to discriminate a novel from a familiar animal. Thus, social novelty and social identity signals are likely conveyed through separate pathways, with the entorhinal cortex providing specific multisensory information about an animal’s identity and novelty detection requiring a local computation within CA2.

## Introduction

Memory formation depends on our ability to detect and distinguish novel from familiar sensory information and then to store that information in long-term memory. The hippocampus, which is classically known for its role in declarative memory, our repository of information of places, objects, events and other individuals, has been found to be important for both novelty detection and long-term memory storage^1–3^. However, the neural circuits by which the hippocampus detects novelty and stores detailed information remain unknown. In particular, it is unclear whether these two functions are mediated by the same or distinct circuits. Here, we address this question by examining the neural circuitry responsible for hippocampal-dependent social novelty recognition and social memory, the ability of an animal to recognize and remember another of its species.

Recently, the CA2 subregion of the dorsal hippocampus has emerged as a critical element of a brain network supporting social recognition memory^4–10^. Although CA2 encodes both social novelty and social identity, the neural circuits that provide these social signals to CA2 are not well understood. One recent study reported that subcortical input to CA2 from the supramammillary nucleus provides information about social novelty^11^. However, it is not known whether this input responds differentially to novel versus familiar conspecifics. Moreover, as the supramammillary nucleus inputs largely target inhibitory neurons in CA2^11^, it is unclear how this might enhance CA2 firing.

In contrast to the net inhibitory influence of the supramammillary nucleus, CA2, like other hippocampal regions, receives its major excitatory input from the entorhinal cortex (EC), a multimodal association area whose medial (MEC) and lateral (LEC) subregions convey spatial and non-spatial information, respectively^12–14^. Information from the EC reaches CA1 and CA2 regions via parallel indirect and direct pathways. In the indirect route, the EC sends excitatory input through the perforant path to the dentate gyrus (DG), whose mossy fibers activate CA3 pyramidal neurons (PNs), which then excite CA1 and CA2 PNs through their Schaffer collaterals. EC also sends direct input to CA1, CA2 and CA3 PNs. However, whereas the direct input only weakly excites CA1 and CA3, this input produces a much stronger activation of CA2 PNs^15,16^.

To date, the relative importance for social memory of the direct versus indirect routes by which information from EC arrives in CA2 is unknown. Moreover, it is not clear whether LEC or MEC inputs are specifically involved in social memory. Finally, we do not know whether these inputs selectively participate in social novelty detection or social memory storage. Here, we use an optogenetic approach to dissect the relative strength of the direct MEC and LEC inputs in exciting CA2 PNs. We then compare the roles of the direct MEC and LEC inputs to CA2 to the indirect inputs that arrive via DG in mediating social memory storage. Finally, we use fiber photometry to ask whether EC inputs to CA2 are activated during a social experience and whether they differentially respond to a novel compared to a familiar animal. Our results indicate that the direct LEC inputs, but not the MEC inputs, are activated during social interaction and provide a strong excitatory drive to CA2 that is required for social memory storage. Moreover, these inputs do not differentiate a novel from a familiar animal, suggesting that social novelty detection in CA2 may be mediated by a distinct input, perhaps from the supramammillary nucleus.

## Results

### Strong dorsal CA2 depolarization by the lateral entorhinal cortex

As a first step in investigating how social information arrives in CA2, we examined the relative contributions of the MEC and LEC direct inputs. Electrical stimulation of MEC and LEC axons with an electrode in the stratum lacunosum moleculare (SLM), the site of the direct inputs, evoked a large postsynaptic potential (PSP) in CA2 PNs in dorsal hippocampal slices (Extended Data Fig. 1a,b), as previously reported^15–17^.

To investigate the differential contribution of the EC regions to the activation of dorsal CA2 neurons, we took advantage of the spatial segregation of the MEC and LEC fiber pathways as they traverse, respectively, the middle and outer molecular layers of DG en route to CA2. The PSP recorded in CA2 PNs evoked by stimulating the LEC pathway was around 1.5- to 2.0-fold larger than the PSP evoked by MEC stimulation (Extended Data Fig. 1c-e). We observed a similar differential response in the size of the extracellular field excitatory postsynaptic potential (fEPSP) recorded in the SLM of CA2 (Extended Data Fig. 2).

To obtain more direct evidence for a preferential role of LEC in exciting CA2 PNs, we used an optogenetic approach to selectively activate MEC or LEC inputs. We injected an AAV into MEC or LEC to express the excitatory light-activated channel channelrhodopsin2 (ChR2). Optogenetic stimulation of these inputs evoked a large PSP in CA2 PNs when ChR2 was expressed in either LEC or MEC. However, similar to the results from electrical stimulation, a significantly greater response was evident with ChR2 expressed in LEC compared to MEC, with LEC-evoked PSPs roughly twice as large as MEC-evoked responses (Fig. 1b-c). This difference was also present when trains of stimuli, rather than single pulses, were used (Fig. 1d). Moreover, the difference between LEC- and MEC-evoked responses was mainly due to differences in excitatory transmission, as we observed a similar relative difference when we blocked inhibition with antagonists of GABA_A_ and GABA_B_ receptors to measure the pure excitatory postsynaptic potential (EPSP) (Extended Data Fig. 3).

**Figure 1.**
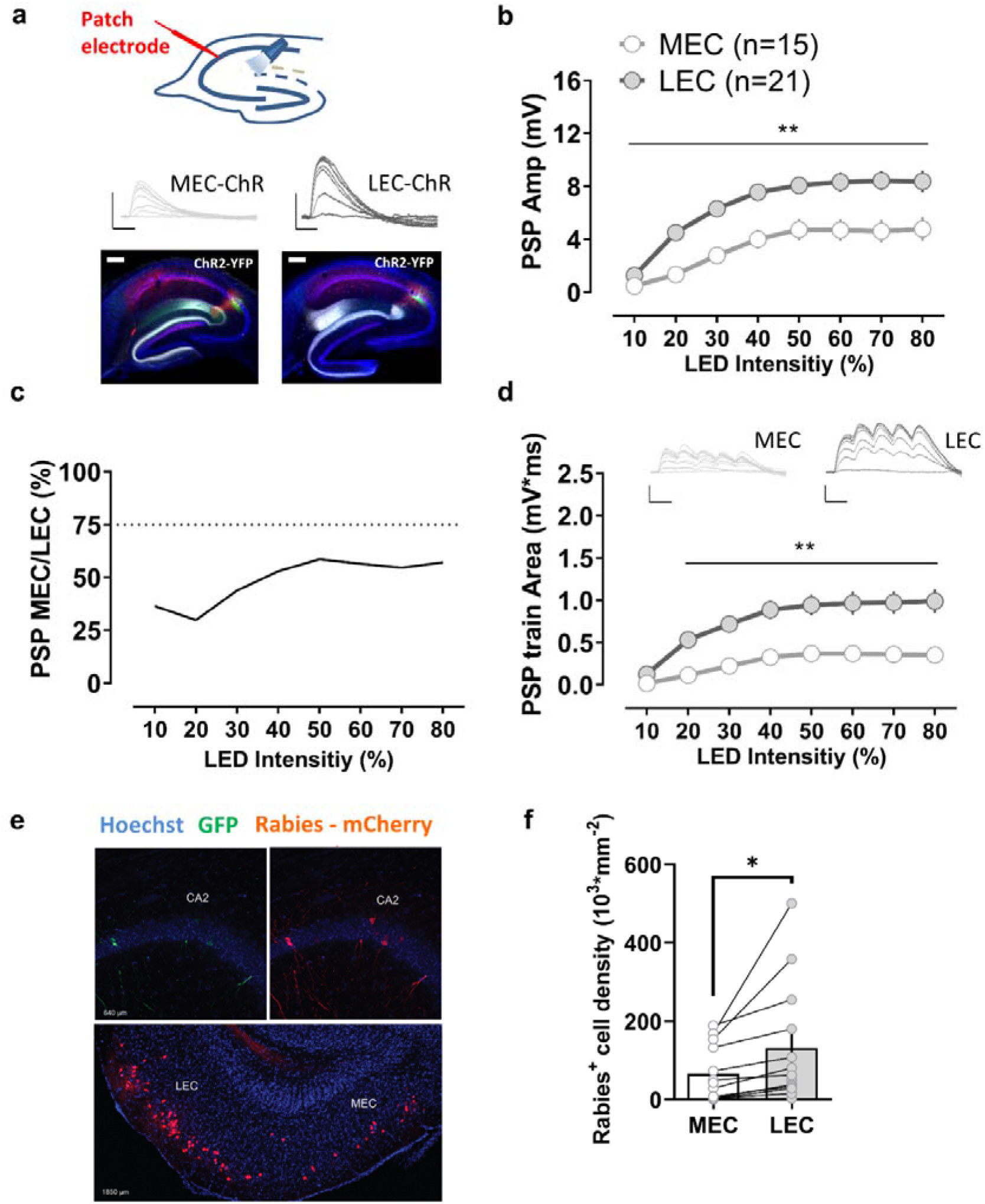
Comparison of synaptic responses of CA2 pyramidal neurons to their direct medial and lateral entorhinal cortical inputs. **a,** An AAV was injected to express ChR2 in the medial (MEC) or the lateral entorhinal cortex (LEC). Photostimulation of ChR2-expressing terminals in the stratum lacunosum moleculare evoked a large postsynaptic potential in CA2 pyramidal neurons in acute hippocampal slices for both, MEC and LEC injected groups. The LEC-evoked response was significantly larger than the MEC-evoked response using either a single light pulse (**b, c**) or a short train of optical stimuli (**d**). **e,** Monosynaptic contacts to dorsal CA2 from the entorhinal area originate largely from the LEC. Retrograde tracing from dorsal CA2 using G-deleted rabies virus expressing mCherry (Rabies-mCherry), after CA2 infection with a Cre-dependent helper virus (expressing GFP). Coronal hippocampal sections in the upper panel and entorhinal horizontal slice in the lower panel. **f,** The number of mCherry positive cells is greater in LEC than MEC. Scale bars: 5 mV/25 ms and 200 μm. *: p<0.05 paired t-test, **: p<0.01 Holm-Sidak’s post hoc test after two-way mixed-design ANOVA.

As a further measure of the relative influence of MEC and LEC inputs, we expressed the inhibitory opsin Archaerhodopsin-T (Arch) in LEC or MEC and examined how selective inhibition of either pathway affected the PSP evoked by electrical stimulation in SLM. Illumination of the SLM with yellow light in slices from mice in which Arch was expressed in MEC or LEC significantly decreased the PSP evoked by electrical stimulation in SLM. However, the extent of inhibition of the PSP was significantly greater when Arch was expressed in LEC than when it was expressed in MEC (Extended Data Fig. 4), supporting the view that the direct LEC input predominates over the MEC input in exciting CA2.

Anatomical support for the larger influence of LEC than MEC on CA2 excitation comes from an examination of the pattern of these inputs when labeled with GFP-tagged ChR2. Consistent with their topology in DG, the two sets of fibers were differentially localized in CA2, with MEC axons localized to a narrow strip in a more proximal domain of SLM in CA2 (closer to the soma) compared to LEC, whose axons occupied a broader more distal region in SLM, extending to the border with DG. Moreover, the width of the LEC projection was ~2-fold larger than that of MEC, consistent with the relative synaptic weights of the two inputs (Extended Data Fig. 5). To gain a more direct measure as to the relative difference in the number of LEC and MEC neurons that project to CA2, we used monosynaptic retrograde tracing by co-injecting G-deleted rabies virus and Cre-dependent helper virus into the CA2 region of *Amigo2-Cre* mice, which express Cre relatively selectively in CA2 PNs. Retrogradely labeled cells were observed in both LEC and MEC, as previously described^4,18^. However, the number of rabies^+^ cells was significantly higher in LEC, roughly twice as great compared to the MEC (Fig. 1e,f).

The above results provide a coherent picture showing that while both MEC and LEC strongly excite CA2 PNs through their direct projections in the dorsal hippocampus, activation of the LEC inputs evokes a CA2 PSP that is 1.5 to 2.0 fold larger than that evoked by MEC inputs. This difference can be accounted for by the roughly 2-fold greater number of LEC neurons that project to CA2 compared to MEC. Next, we asked whether these differences in synaptic strength were reflected in the behavioral influence of these two regions on CA2-dependent social memory.

### Disrupting the lateral entorhinal input to dorsal CA2 impairs social memory

To investigate the role of the EC projections to dorsal CA2 in social memory, we expressed Arch or GFP in LEC or MEC and illuminated their projections in dorsal CA2 with yellow light as animals were engaged in an open arena, two-choice social memory task^6^. In the learning phase of this task, a subject mouse is allowed to explore for 5 min a square arena in which two novel stimulus mice (S1 and S2) are presented in wire cup cages in opposite corners of the arena. The subject mouse is then placed in its home cage for 30 min, after which a memory recall test is performed, in which the subject mouse explores for 5 min the same arena, in which one of the two original stimulus mice is replaced with a third novel mouse (N). Social memory was assessed by the preference of the subject mouse to explore the novel mouse (N) compared to the original, now-familiar stimulus mouse (S1 or S2). We quantified this preference with a discrimination index (DI) = (time exploring mouse N – time exploring mouse S) / (time exploring mouse N + time exploring mouse S).

We first examined the effects of silencing the LEC inputs to CA2. As expected, control mice expressing GFP in LEC spent significantly more time interacting with a novel animal compared to the now-familiar mouse in the memory recall trial (Fig. 2b,e). Furthermore, control mice spent significantly less time exploring the now-familiar mouse in the recall trial than they spent exploring the same mouse during the learning trial (Fig. 2d,g). Illumination of the CA2 region with yellow light during either the learning or recall trials did not affect memory performance in the control group (Fig. 2). In contrast, in mice expressing Arch in LEC, illumination of dorsal CA2 with yellow light during either the learning or recall trials produced a significant impairment in social memory performance (Fig. 2). Thus, after applying light during either the learning or recall trials, animals expressing Arch in LEC spent a similar amount of time exploring the novel and the now-familiar mouse during the recall trial (Fig. 2b,e), suggesting an impairment in distinguishing between novel and familiar conspecifics. Moreover, Arch-expressing mice did not show any significant reduction in the time spent exploring the now-familiar mouse during the recall trial (Fig. 2d,g) when yellow light was applied during either the learning or recall trial, confirming a deficit in recognition memory.

**Figure 2.**
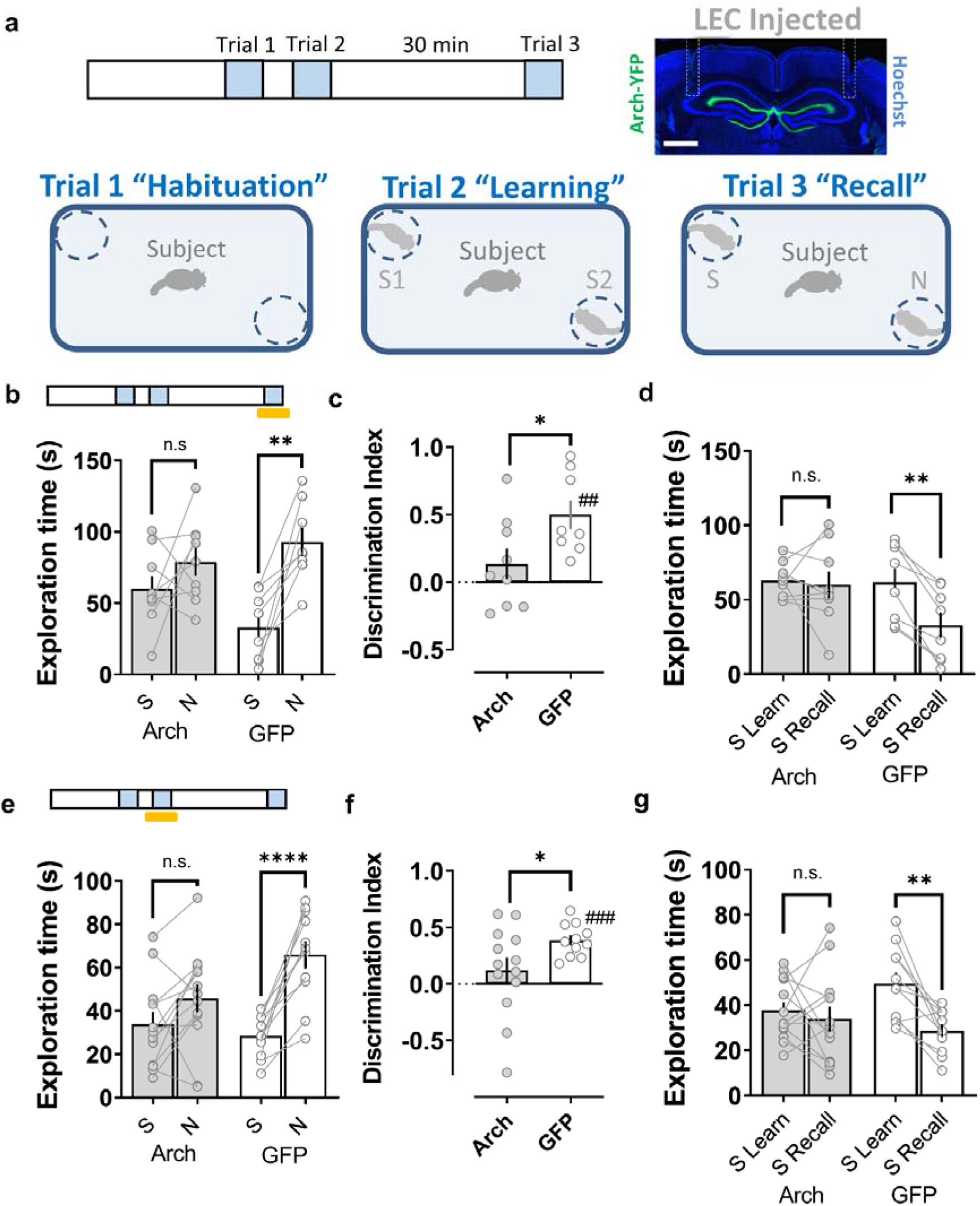
Disrupting the lateral entorhinal cortical input to dorsal CA2 impairs social memory. **a**, Schema of the two-choice social memory task consisting of three trials. Mice first explored the arena with two empty cups (trial 1 habituation). Next, two novel stimulus mice (S1 and S2) were placed one in each cup; the subject mouse was allowed to explore the arena with the two novel mice for 5 min in trial 2 (learning). The subject mouse was then removed from the area for a 30min inter-trail interval, after which it was reintroduced to the arena in a 5 min recall trial (trial 3), in which one of the stimulus mice presented in trial 2 (S1 or S2) was replaced by a third novel mouse (N). Social memory is manifest as: 1. A greater time spent exploring mouse N compared to the now familiar stimulus mouse (S1 or S2) in the recall trial (**b, e**) as quatified by a discrimination index (**c, f**) and 2. A decreased time spent exploring S1 or S2 in the recall trial relative to the learning trial (**d, g**). Inset shows the expression of Arch in the lateral perforant path and the optical fiber location (dashed outline) in a coronal brain slice from a mouse previously injected in lateral entorhinal cortex (LEC) with an Arch-expressing AAV. Shining yellow light on LEC inputs in dorsal CA2 during the recall phase of the task (trial 3) (**b-d**) or during the learning phase (trial 2) (**e-g**), impairs social memory performance of animals expressing Arch in LEC relative to the control group expressing GFP. Scale bar: **a**: 1 mm. *: p<0.05 t-test. ##: p<0.01, ###: p<0.001 one-sample t-test against “0”. **: p<0.01, ****: p<0.0001 Holm-Sidak’s post hoc test after two-way mixed-design ANOVA.

In contrast to the deficits in social memory observed when the LEC inputs to CA2 are silenced, optogenetic silencing of the MEC inputs to CA2 did not inhibit social memory, either when illumination was applied during the learning or recall trials (Fig. 3). Thus, mice expressing Arch in MEC showed a normal preference for the novel animal in the recall trial, regardless of whether light was applied during the learning or recall trial. Moreover, illumination of CA2 of Arch-expressing mice did not prevent the normal decreased exploration of the familiar mouse in the recall trial relative to the exploration of the same mouse in the learning trial. Indeed, rather than suppressing social memory, there was a trend for social discrimination to be enhanced upon silencing of MEC inputs to CA2, although this enhanced performance was not significant.

**Figure 3.**
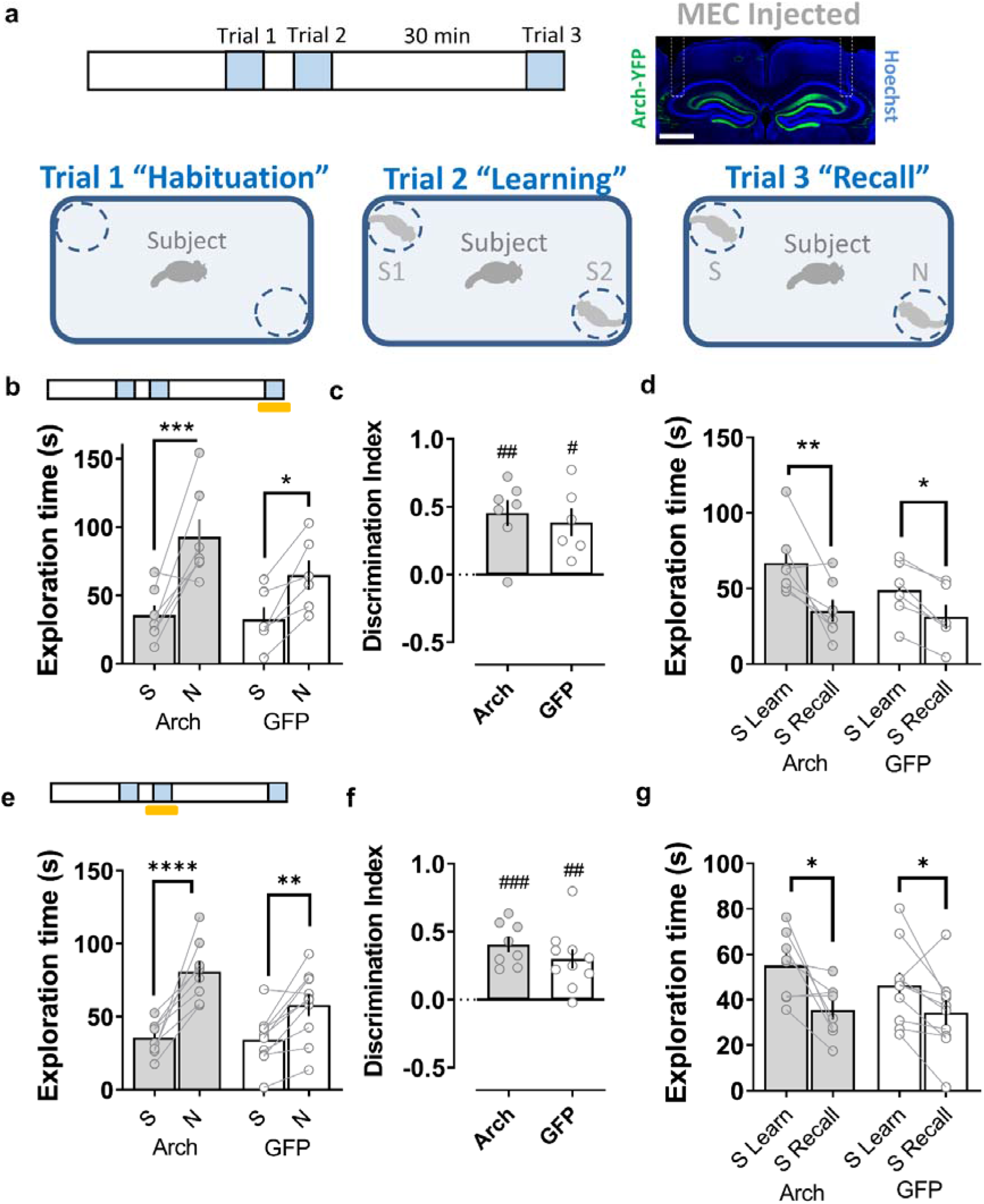
Disrupting the medial entorhinal cortical input to dorsal CA2 does not affect social memory. **a**, Schema of the social memory task described in Figure 2. Inset shows the expression of Arch in the medial perforant path in a coronal brain slice from a mouse previously injected in the medial entorhinal cortex (MEC) with an Arch-expressing AAV. Shining yellow light on MEC inputs in dorsal CA2 during the recall phase of the task (trial 3) (**b-d**) or during the learning phase (trial 2) (**e-g**) does not impair social memory in mice expressing Arch in MEC relative to the control group expressing GFP. Scale bar: **a**: 1 mm. #: p<0.05, ##: p<0.01, ###: p<0.001 one-sample t-test against “0”. *: p<0.05, **: p<0.01, ***: p<0.001, ****: p<0.0001 Holm-Sidak’s post hoc test after two-way mixed-design ANOVA.

The social memory deficits seen with optogenetic silencing of the LEC inputs to CA2 were not attributable to a general impairment of novelty detection as optogenetic silencing of LEC terminals in CA2 did not affect performance in an analogous novel object recognition test (Extended Data Fig. 6). Moreover, the lack of social recognition, which requires recognition of olfactory cues, was not due to a general impairment in olfaction as optogenetic silencing of either LEC or MEC inputs to CA2 did not impair an animal’s ability to find a hidden food pellet (Extended Data Fig. 7). Thus our data suggest that the information that LEC, but not MEC, provides to CA2 plays a specific role in social novelty discrimination.

To explore the generality of our behavioral results, we investigated how disrupting the communication between LEC or MEC and dorsal CA2 affected social recognition using a different behavioral paradigm, the direct interaction task (Extended Data Fig. 8-9). In the direct interaction task, the subject mice are allowed to freely explore a novel juvenile male stimulus mouse for 2 min during a learning trial. The stimulus mouse is then removed from the arena and after a 30 min inter-trial interval it is reintroduced into the test arena for a recall trial. In this task, social memory is manifest as a reduction in the exploration time of the same juvenile in the recall trial relative to the learning trial. Control mice expressing GFP in LEC terminals showed a significant reduction in exploration time when the same juvenile was reintroduced in the recall trial, independently of whether dorsal CA2 was illuminated with yellow light during the learning or recall trials. In contrast, mice expressing Arch in LEC that were illuminated with yellow light during either the learning or recall trials failed to show a decrease in the exploration of the juvenile in the recall trial (Extended Data Fig. 8). Optogenetic silencing of MEC inputs to CA2 during either the learning or recall trials caused no impairment in social memory (Extended Data Fig. 9). As a control to rule out the possibility that decreased exploration time during the recall trial resulted from fatigue of the subject mice or lack of motivation for social exploration, rather than a manifestation of social memory, we introduced a second novel juvenile mouse during the recall trial. In all experimental groups, subject mice explored the two novel mice to equal extents in the two trials, indicating that the decreased exploration of the familiar mouse in a recall trial did indeed represent social memory (Extended Data Fig. 10).

### Information flow through the dorsal trisynaptic path is dispensable for social memory

Although our results above suggest that the direct inputs from LEC to CA2 are required for social memory, it is also possible that information conveyed through the trisynaptic path, from EC to DG to CA3 to CA2, or through a disynaptic path, from EC to DG to CA2^18^, is also necessary. Furthermore, our approach using a fiber optic probe targeting EC axons in CA2 may have allowed sufficient light to reach the EC axons in dorsal DG, thereby inhibiting the activation of DG by its EC inputs. Thus, to explore any possible contribution of DG activation to the behavioral effects we observed with optogenetic silencing of the EC inputs in CA2, we directly examined the effect of silencing DG granule cells on social memory performance.

To ensure that we silenced as large a population of DG granule cells as possible, we examined mice in which a DG granule cell Cre driver line (*POMC-Cre*)^19^ was crossed with a mouse line expressing the inhibitory DREADD hM4Di under control of Cre^20^. Delivery of the DREADD agonist CNO through an implanted cannula to dorsal DG was sufficient to effectively reduce DG activity in these mice. Thus, CNO infusion 30 min before an i.p. injection of the convulsant pilocarpine resulted in a significantly lower number of c-Fos^+^ cells compared to a Cre-control group that had also been infused with CNO (Fig. 4a).

**Figure 4.**
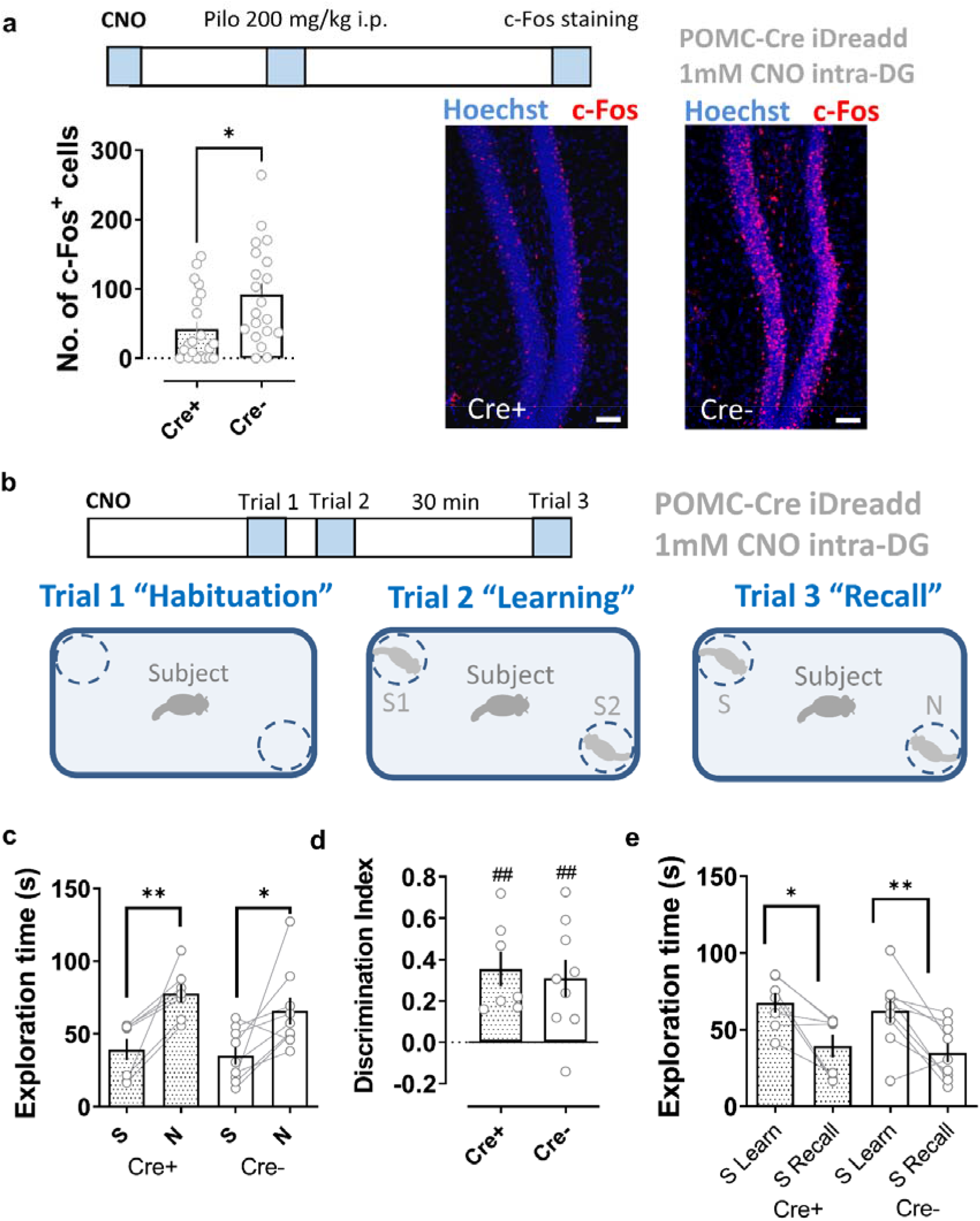
Pharmacogenetic silencing dorsal dentate gyrus granule cells does not impair social memory. **a,** A pharmacogenetics approach effectively reduces dentate granule cell activity. The inhibitory DREADD (iDREADD) agonist CNO (1 mM, 1 μl per side) was infused into the dorsal dentate gyrus of *POMC-Cre+* x Rosa26-LSL-hM4Di-mCitrine (Cre+) mice expressing iDREADD in DG granule cells or control mice (*POMC-Cre*- x Rosa26-LSL-hM4Di-mCitrine, Cre-). After 30 min mice were given an i.p. injection of pilocarpine. CNO significantly reduced the number of c-Fos+ neurons in the iDreadd (Cre+) group compared to that of controls (Cre-). **b,** Schema of the social memory task. **c-e,** CNO injection into the dentate gyrus of iDreadd (Cre+) mice did not significantly impair social memory performance compared to the control group (Cre-). In **a** Scale bar: 50 μm. *: p<0.05, t-test. In **c-e** ##: p<0.01 one-sample t-test against “0”. *: p<0.05, **: p<0.01, Holm-Sidak’s post hoc test after two-way mixed-design ANOVA.

In contrast to the ability of CNO to inhibit DG activity, infusion of the DREADD agonist into DG of mice 30 min prior to the two-choice social recognition test did not impair social memory. Thus CNO-treated Cre+ mice and Cre-control animals both explored the novel mouse for a significantly greater time than the familiar mouse during the memory recall trial (Fig. 4c,d). Furthermore, both groups of mice spent significantly less time exploring the now-familiar mouse in the recall trial than during the learning trial (Fig. 4e). Our findings thus show that dorsal DG granule cells do not significantly contribute to social memory in mice, implying that it is indeed the direct LEC inputs to CA2 that are of key importance.

### Input from the lateral entorhinal cortex to CA2 is selectively enhanced during social exploration

How does the LEC participate in social memory and what is the basis for its selective involvement in social memory relative to MEC? To gain insight into these questions we investigated whether LEC or MEC responds to socially-relevant information by staining these regions for expression of the immediate-early gene c-Fos as a marker of neuronal activity following the recall phase of the two-choice social memory task. We observed a significant increase in the number of c-Fos^+^ cells in both MEC and LEC following social exploration, relative to c-Fos^+^ cells in mice kept in their home cage (Fig. 5). Although the overall number of c-Fos^+^ cells was similar in LEC and MEC, we saw significant differences when we classified cells as to whether they were present in the superficial layers of EC, whose cells project to hippocampus, or deep layers of EC, whose cells receive input from CA1. In LEC we observed significantly more c-Fos+ cells in superficial compared to deep layers (p<0.001, paired t-test), whereas in MEC the two layers contained similar numbers of c-Fos^+^ cells. Furthermore, the number of c-Fos^+^ cells in the superficial layers of LEC was significantly greater than the number of c-Fos^+^ cells in superficial layers of MEC (p<0.01, paired t-test).

**Figure 5.**
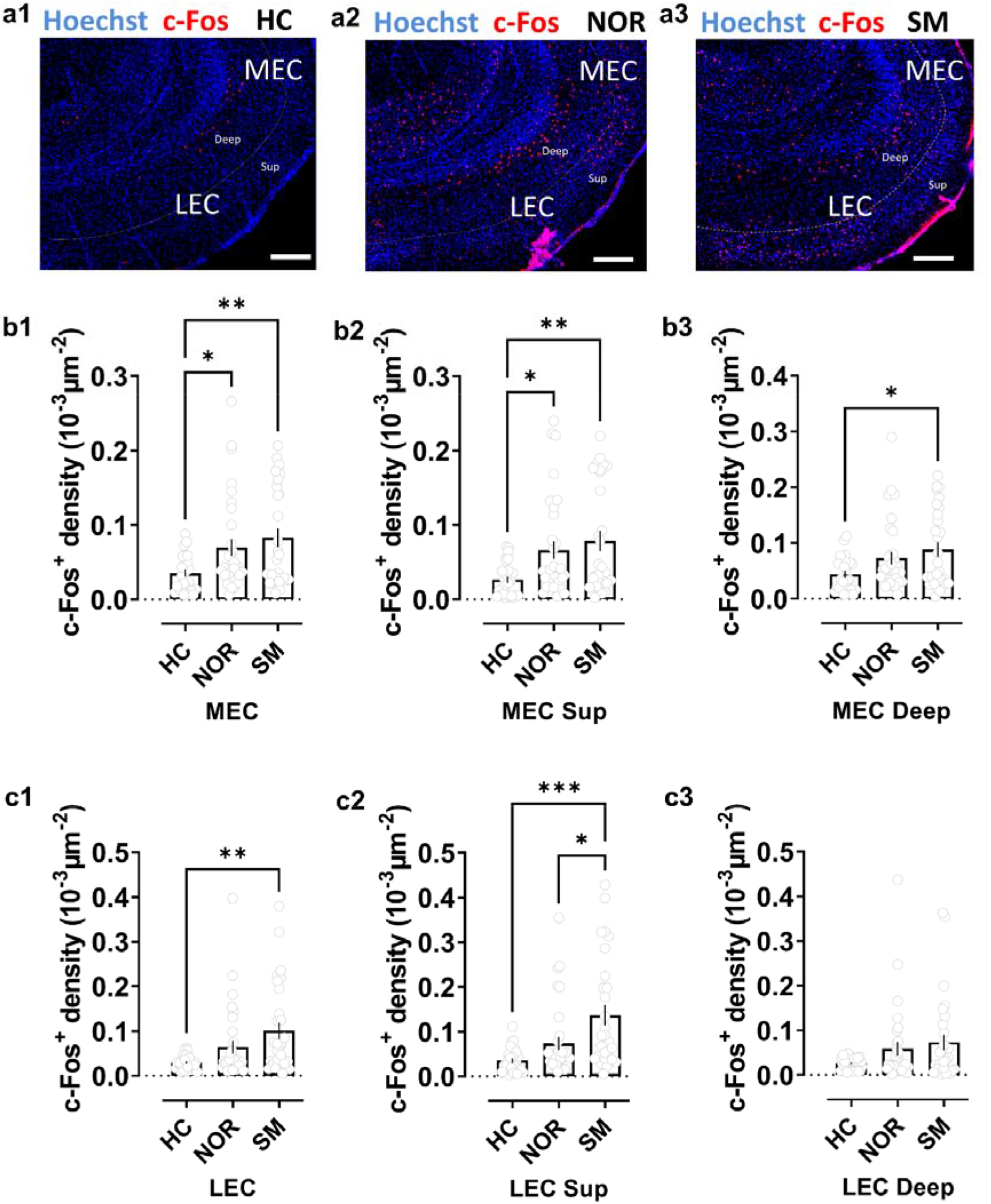
Social exploration preferentially activates lateral compared to medial entorhinal cortex. **a**, Staining for c-Fos in lateral (LEC) and medial (MEC) entorinal cortices in horizontal brain slices from mice in control home-cage conditions (HC) or following novel object recognition (NOR) or social memory (SM) tasks. **b**, Quantification of c-Fos+ cell density in MEC in both deep and superficial layers combined (**b1**) or separated into superficial (**b2**) or deep layers (**b3**). Mice subjected to the NOR or SM task showed a significantly larger density of c-Fos+ cells than mice in HC conditions in both entire MEC and in superficial MEC layers. In deep MEC layers the increase over HC c-Fos+ staining was only significant following the SM task, although there was a trend in the NOR task. There was no significant difference in c-Fos+ density following SM compared to NOR task in total MEC (**b1**) or individual layers (**b2, b3**) **c**, In superficial and deep LEC layers combined (**c1**) and superficial LEC layers alone (**c2**), we observed a significant increase in c-Fos+ density compared to HC levels following the SM task, with no significant increase following the NOR task (although there was a trend). We saw no significant change in either SM or NOR tasks in deep LEC alone (**c3**). c-Fos+ density in superficial LEC was significantly greater following SM task compared to NOR task. Scale bar: 200 μm. *: p<0.05, **: p<0.01, ***: p<0.001 Holm-Sidak’s post hoc test after one-way ANOVA.

To determine whether the increase in c-Fos+ cells was specific for social novelty, we examined mice that had performed a two-choice novel object recognition task, analogous to the social recognition task. Although we found a significant increase in c-Fos^+^ cells in MEC following exploration of a novel object, relative to animals in their home cage, there was no significant increase in c-Fos^+^ cells in LEC (Fig. 5). Moreover, the number of c-Fos^+^ cells in the superficial layers of LEC was significantly greater following social compared to object exploration. These results suggest that LEC is selectively tuned to respond to social signals, relative to MEC, consistent with its behavioral importance for social memory.

Finally, we assessed the dynamics of LEC input activity in dorsal CA2 during social and object exploration using fiber photometry measurements of Ca^2+^ levels. We expressed the genetically encoded fluorescent Ca^2+^ sensor GCaMP7f in LEC using targeted viral injections. We then implanted an optic fiber in the SLM region of dorsal CA2 and measured Ca^2+^ levels in the population of LEC inputs to CA2 based on GCaMP7f fluorescence (Fig. 6a). Bouts of exploration of a novel conspecific elicited a large, consistent increase in GCaMP7f fluorescence. In contrast, object exploration produced a weak, non-significant increase in fluorescence, consistent with the c-Fos staining patterns observed above. We then repeated the Ca^2+^ measurements during social exploration of a familiar littermate. Despite the lack of social novelty, the littermate elicited large increases in GCaMP7f fluorescence that were comparable to the levels seen during exploration of a novel animal (Fig. 6b,c). These results thus suggest that LEC, which is critical for social memory, conveys similar overall levels of synaptic input to the CA2 region in the dorsal hippocampus regardless as to whether an animal explores a novel or familiar conspecific.

**Figure 6.**
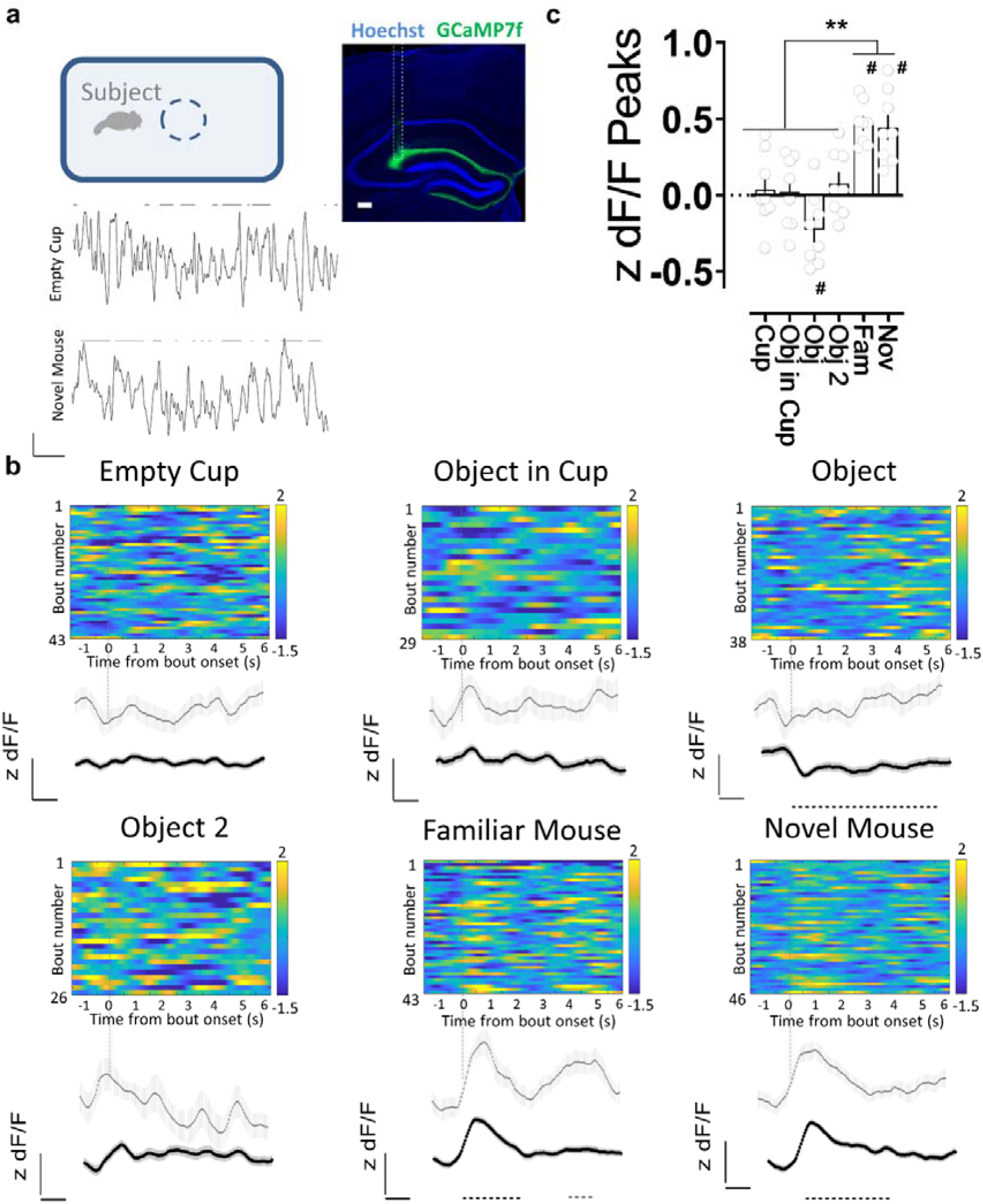
Activity of direct lateral entorhinal cortex input to dorsal CA2 increases during social exploration. **a**, Fiber photometry recordings of GCaMP7f fluorescence in lateral entorhinal cortex (LEC) inputs to CA2 in dorsal hippocampus while an animal explores different items in an open arena. Coronal section of the hippocampus showing the expression of GCaMP7f and the optical fiber location (dashed outline) in mice previously injected with AAV in LEC. GCaMP7f z-scored dF/F traces during bouts of interaction (lines at top) with a novel object (empty cup) or a novel mouse. **b**, Exploration of a littermate or a novel conspecific, but not novel objects, is associated with an increase in GCaMP7f fluorescence intensity in LEC inputs in dorsal CA2. Color-coded z-scored dF/F traces from a single animal aligned to the time of interaction. Gray trace shows average fluorescence from all interaction bouts of a given type for that animal. Black traces show average of all animals (n=8). **c**, Mean peak fluorescence values for given bouts of exploration averaged from all animals. Scale bars: **a**: inset 200 μm and traces 1z unit dF/F, 10 s **c**: 0.5 z units dF/F, 1 s. Color scale, number indicate range of z scores. #: p<0.0001 one-sample t-test against “0”. **: p< 0.01, Holm-Sidak’s post hoc test after one-way ANOVA.

## Discussion

Although the medial and lateral divisions of EC are well known to provide, respectively, distinct spatial and non-spatial forms of information to the hippocampus^13,14^, there has been no systematic study of the relative roles of these cortical subdivisions and their routes of information flow in hippocampal-dependent social memory. Furthermore, although there have been several proposed models for the function of convergent direct and indirect cortical inputs to CA1, CA2 and CA3 regions, including the detection of novelty^21^ or salience^22^, memory specificity^23^, or prevention of memory interference^24^, there have been relatively few experimental tests of such proposals. Here we provide the first direct evidence that the direct inputs from LEC to CA2 are critical for both the encoding and recall of social memory. Moreover, our data suggest that the detection of social novelty is not provided by LEC inputs but depends on a local computation in CA2.

Compared to our detailed understanding of how MEC encodes spatial information through grid cell, border cell and head direction cell activity^25,26^, we know much less about how representations encoded in the LEC contribute to hippocampal-dependent memory. Recent work in animal models and humans suggests that LEC might process temporal memories^27,28^, egocentric information^29^ and episodic-like representations^30^. Also, input from LEC to the hippocampus has been found to be required for olfactory associational learning^31^. This emerging picture points to more varied and higher-order representations in LEC than the more selective spatial information conveyed by MEC. Indeed, LEC constitutes a central cortical hub, forming one of the richest sets of association connections of any brain region^32,33^. Our present results extend these findings by showing that the encoding and recall of social memory require the direct input of multisensory information from LEC, but not MEC, to the dorsal CA2 region of the hippocampus.

Using a combination of ex vivo electrophysiology, optogenetic and chemogenetic behavioral studies, and fiber photometry, we explored the relative roles of the direct LEC and MEC inputs in the synaptic excitation of CA2 PNs, social memory behavior and the encoding of social exploration. At the physiological level, we find that LEC provides a significantly stronger direct excitatory drive to CA2 PNs compared to MEC. The preferential excitation of CA2 by LEC compared to MEC is reflected in our behavioral results showing that LEC but not MEC inputs to dorsal CA2 are required for CA2-dependent social memory. The selective behavioral role of LEC compared to MEC in social memory is consistent with our findings that LEC is more strongly and selectively activated by social exploration compared to MEC, as revealed by c-Fos labeling.

In addition to defining the relative importance of LEC and MEC in social memory, our study also provides evidence that the direct LEC inputs to CA2 are more critical for social memory compared to the indirect EC inputs, which reach CA2 via the DG and CA3 regions of the hippocampus. Although a prior study suggested that the EC inputs to DG were also important for social memory^34^, the experimental approaches employed in that study could not distinguish whether DG or some other downstream target of the EC inputs, such as CA2, was involved. Our finding that direct silencing of dorsal DG fails to suppress social memory is consistent with the view that the EC inputs participate in social memory predominantly through the direct excitation of CA2. Although our study cannot definitively rule out the possibility that the direct EC inputs to CA3 may also participate in social memory, the EC inputs provide only weak excitation of CA3 compared to CA2^35^. Moreover, another study found that silencing dorsal CA3, and to some extent dorsal DG, through a chemogenetic approach cause no significant impairment in social memory^36^.

Besides the classic division of hippocampus into its DG, CA3, CA2 and CA1 regions along its transverse axis, there is a well-known heterogeneity in anatomical connectivity, function and behavior along the longitudinal, dorsal-ventral (or septal-temporal) axis of the hippocampus^2^. Our study here focused on the projections of EC to the dorsal region of CA2, as this portion of CA2 is critical for social memory^4,5^. In contrast to the importance of dorsal CA2, neither dorsal CA1^37^ nor dorsal CA3^36^ appear to be required for social memory. Rather, the ventral portions of both CA1^37^ and CA3^36^ have been found to play important roles in social memory. The apparent dichotomy between the importance of dorsal CA2 versus ventral CA1/CA3 was resolved by Meira et al. (2018), who found that dorsal CA2 participates in social memory through its longitudinal excitatory projections to the ventral hippocampus. The extent to which ventral CA2 participates in social memory and the importance of the direct entorhinal inputs to ventral hippocampus remain to be determined.

Our study also provides insight into the functional role of the direct and indirect cortico-hippocampal circuit architecture. As noted above, one interesting model posits that this circuitry is important for the ability of the hippocampus to serve as a novelty detector^21^, with the direct inputs providing an immediate representation of sensory experience that is then compared with mnemonic information from stored representations in DG and CA3 conveyed by the indirect inputs to either CA1 or CA2. In this manner, the hippocampus can gauge both familiarity, the ability to distinguish a novel from a previously encountered stimulus, and store and recall the detailed sensory information that a given experience comprises^38^.

In vivo recordings have previously shown that a significant subset of CA2 PNs fire preferentially during interactions with a novel animal compared to a familiar littermate^7^. Moreover, CA2 population firing rates can be used to train a linear decoder to distinguish whether an animal is interacting with a novel or familiar animal, indicating that CA2 encodes social novelty^7^. Our finding that the global activity conveyed by LEC to CA2, assessed by mean population Ca^2+^ levels measured through fiber photometry, are identical for novel and familiar animals suggests that the LEC itself is unlikely to be the source of the novelty signal, implying that the determination of social novelty is likely to be computed locally in CA2.

One potential source of novelty information is through the inputs CA2 receives from the hypothalamic supramammillary nucleus, which has been recently shown to convey social novelty signals to CA2^11^. However, it is not yet known whether the supramammillary inputs differentiate between a novel versus a familiar animal. Moreover, because the supramammillary inputs preferentially excite inhibitory neurons in the CA2 region^11,39^, it is unclear how such an input could enhance CA2 PN firing to social novelty. Finally, as the computation of novelty requires processing of complex sensory information and its comparison to stored representations, such computations are likely performed in higher-order brain regions.

Here we propose a model for the computation of social novelty based on the intrinsic circuitry of the hippocampus and the finding that information relayed to CA2 from DG and CA3 through the trisynaptic path elicits strong net feedforward inhibition of CA2 pyramidal neurons^15^. According to this view, the specific set of sensory cues that constitute the unique identity of a given novel or familiar conspecific would produce a strong activation of CA2 PNs due to the strong excitation they receive from the direct LEC inputs. However, for interactions with a familiar conspecific, activation of the stored representations of that conspecific in DG and/or CA3 would produce feedforward inhibition of CA2, resulting in an enhanced response to a novel social stimulus. Although this possibility seems at odds with our finding that silencing dorsal DG fails to affect social memory or with a study showing that silencing dorsal CA3 also does not alter social memory^36^, it is possible that longitudinal projections to CA2 from more ventral regions of hippocampus that are known to participate in social memory may provide the relevant mnemonic information.

As social interactions are at the core of everyday experience, and socially-related psychiatric disorders are a serious mental health problem, understanding the interplay of brain structures supporting adaptive social behavior is of key importance. Indeed, the *Df(16)A^+/-^* mouse line, a genetic model of the human 22q11.2 microdeletion, which is one of the greatest known genetic risk factors for schizophrenia^40^, has impaired social memory^41^ that is associated with impaired CA2 firing responses to social cues and social novelty^7^. Interestingly, these mice also have a decrease in CA2 feedforward inhibition due to the loss of parvalbumin-positive interneurons^41^, whose loss is also seen in the general population of individuals with schizophrenia and bipolar disorder^42,43^. Such a decrease in feedforward inhibition due to trisynaptic inputs could contribute to the impaired social novelty detection. Furthermore, neurons in LEC superficial layers are particularly susceptible to damage in early Alzheimer’s disease stages^44–46^, which could also contribute to abnormal social memory associated with this disorder. Thus, elucidating fundamental questions concerning the role of LEC in physiological conditions might provide insights into better preventive and palliative treatments in pathological neuropsychiatric and neurodegenerative contexts.

## Methods

### Experimental models

All animal procedures were performed in accordance with the regulations of the Columbia University IACUC. 8- to 12-week-old male mice were used for most experiments. The mice were group housed and maintained in a temperature- and humidity-controlled room on a 12:12 h light/dark cycle. All animals were provided with food and water ad libitum. All tests were conducted during the light cycle.

Wild-type C57BL/6J male mice were obtained from the Jackson Laboratory. POMC-Cre(+/-) male mice were obtained from Jackson Laboratory and were bred with R26-hM4Di/mCitrine female homozygous.

### Viral injections

Viral injections were performed as described previously^47,48^. Briefly, mice were anesthetized with isoflurane and placed in a stereotaxic apparatus. A craniotomy was performed above the target region and a glass micropipette was used for viral injection. Injections were performed using a nano-inject II (Drummond Scientific). Twenty-three nl of solution were delivered every 15 s until the total amount was reached. The micropipette was retracted after 5 min. We bilaterally injected 368 nl of AAV2/9 hSyn.hChR2(H134R).eYFP.WPRE.hGH (UPenn Vector Core) or AAV2/9 CaMKII.ArchT-GFP (UNC Vector Core) or pGP-AAV-syn-jGCaMP7f-WPRE (Addgene) to the LEC or MEC. The positions were: −3.4 mm AP, +/- 4.7 mm ML and 2.8 mm DV for LEC injections and −4.9 mm, +/- 3.4 mm ML and 2.8 mm DV for MEC injections. Mice were allowed to recover for 2-3 weeks.

#### Rabies and AAV helper virus injection in CA2

We delivered 50 nl of a G-deleted rabies helper virus AAV2/8 syn.DIO.TVA.2A.GFP.2A.B19G (UNC vector core) into the dorsal hippocampus of Amigo2-Cre mouse at the following coordinates AP −1.8 mm, ML +2.5 mm, DV −1.7 mm. Following 2 weeks of recovery and AAV expression, a second surgery was performed and 300 nl of rabies SAD.B19.EnvA.ΔG.mCherry (SAD-B19 strain, Addgene Cat# 32636 prepared by the Salk institute vector core) was injected at the same coordinates. Mice were killed 7 d later and the brains cut horizontally for entorhinal cortex imaging or coronally for hippocampus imaging.

### Optical fiber implantation

Multimode fibers of 200 μm core and 0.39 numerical aperture (Thorlabs) were used for behavior experiments. The fibers were glued to a ceramic ferrule and polished to enhance coupling efficiency. The optical fibers were implanted in the dorsal CA2 area (−2.0 mm AP, +/- 2.2 mm ML and 2.0 mm DV) 2 weeks after viral injection and fixed to the skull with dental cement. Fibers were coupled to an external fiber using standard FC connectors via a mating sleeve connected to a 589-nm laser (Laserglow).

### Cannula guide implantation

Mice were implanted with a cannula guide extending for 1 mm (Plastics One) below the pedestal. The scalp was removed and holes were drilled (−2.0 mm AP, +/- 1.0 mm ML). Cannula guides were kept in place using super-glue and dental cement. Dummy cannulas (Plastics One) were inserted into the guides. For CNO infusion, mice were placed under light isoflurane anaesthesia and the dummy cannula was removed. The injector cannula protruding 1.5 mm from the cannula guide was then inserted. One microlitre of a 1 mM CNO solution was infused over 2 min. The injector cannula was removed 2 min after the end of the micro-infusion to avoid pulling out the drug and the dummy cannula was put back. Behavioural testing started 30 min after drug infusion.

### Immunohistochemistry

Mice were anesthetized, and brains were processed as previously described^48^. Briefly, after fixation in 4% PFA overnight floating sections were prepared and rinsed three times in 1x PBS and then blocked in 1x PBS with 0.5% Triton X-100 and 5% goat serum for 2 hr at room temperature (RT). Incubation with primary antibodies was performed at 4°C overnight in 1 x PBS with 0.5% Triton X-100. Sections were then washed three times in 1x PBS and incubated with secondary antibodies for 2 hr at RT. Hoechst counterstain was applied (Hoechst 33342 at 1:1000 for 30 min in PBS at RT) prior to mounting the slice using fluoromount (Sigma-Aldricht).

For post-hoc immunocytochemistry after patch-clamp recordings, slices were fixed for 1 h in PBS with 4% PFA and Streptavidin conjugated to Alexa 647 (1:500, ThermoFisher Scientific) was applied during secondary incubation following blocking and permeabilization.

For c-Fos labelling, the first incubation was performed with rabbit anti-c-Fos (1:5000, SySy, 226 003) at 4 °C overnight. The secondary incubation was performed with and anti-rabbit conjugated to Alexa 647 (1:500, ThermoFisher A31573)

### In vitro electrophysiology

Male mice 7-9 weeks old were anesthetized and killed by decapitation in accordance with institutional regulations, as previously described^16,35^. Hippocampi were dissected out and transverse slices from the dorsal hippocampus were cut with a vibratome (Leica VT1200S, Germany) on in ice-cold dissection solution containing (in mM): 125 NaCl, 2.5 KCl, 20 glucose, 25 NaHCO_3_, 1.25 NaH_2_PO_4_, 2 Na-Pyruvate, 2 CaCl_2_ and 1 MgCl_2_, equilibrated with 95% O_2_/5% CO_2_ (pH 7.4). The slices were then incubated at 33°C for 25 min and then kept at room temperature for at least 1 hr before transfer to the recording chamber. All electrophysiological recordings were performed at 31-32°C.

Patch pipettes were pulled from a horizontal micropipette puller (Sutter) and filled with an intracellular solution containing the following (in mM): 135 K-Gluconate, 5 KCl, 0.1 EGTA-Na, 10 HEPES, 2 NaCl, 5 ATP, 0.4 GTP, 10 phosphocreatine. The pH was adjusted to 7.3 and the osmolarity to 290 mOsm. Pipettes of a 3-5 MΩ tip resistance were used. Whole-cell “blind” patch-clamp configuration was established, and cells were held at −70 to −73 mV.

### Two-choice social memory test

This test was performed as previously described^6^. In brief, a subject mouse was habituated for 5 min to a rectangular arena with two empty wire cups in opposite sides. After this, in a learning trial, a novel stimulus male mouse that had no prior contact with the subject mouse was placed inside each of the cups and the subject mouse was allowed to explore the arena with the two novel mice for 5 min. The subject mouse was then isolated for 30 min and one of the stimulus mice was exchanged for a third novel mouse. In the “recall” phase of the test the subject mice were exposed to one of the now-familiar stimulus mice, previously encountered during the learning trial, along with the novel stimulus mouse. Social exploration was quantified as the time spent in active exploration within 5 cm of the perimeter of the cup. We then assessed social memory using a discrimination index:

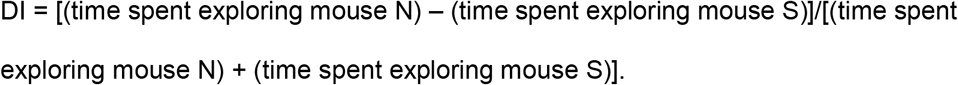

### Direct interaction test

This test was adapted from Kogan et al^49^. Subject mice were placed in a standard clean cage for a 30 min habituation session immediately prior to the experimental sessions. In trial 1, a novel male juvenile stimulus mouse around 5-weeks-old was then introduced into the cage and activity was monitored for 2 min and scored online for social exploration (sniffing, following and allogrooming) initiated by the test subject. The stimulus mouse was then removed from the cage. In trial 2, after an inter-trail interval of 30 min, the subject mouse was allowed to interact for another 2 min with either the previously encountered stimulus mouse or a novel stimulus mouse. Social memory is normally manifest as the decreased exploration of the same stimulus mouse in trial 2 relative to trial 1. In contrast there is normally no decrease in exploration time when the novel mouse is introduced in trial 2, demonstrating that the decreased exploration of the same mouse in trial 2 is not due to fatigue or loss of motivation over the test duration.

### Buried food test

The mice were food-deprived for 18 h before the test, to improve sensitivity. A pellet of the same chow the animals were regularly fed with was hidden under 1 cm of standard cage bedding. The subject mouse was placed in the cage, and the latency to find the pellet was recorded^50,51^.

### Novel object recognition test

A subject mouse was habituated for 5 min to a rectangular arena. After this, two novel objects were placed in opposite sides of the arena and the subject mice were allowed to explore for 10 min (learning phase). After 30 min, one of the objects was exchanged for another novel one and the subject mice were allowed to explore for 5 min. We then assessed memory using a discrimination index:

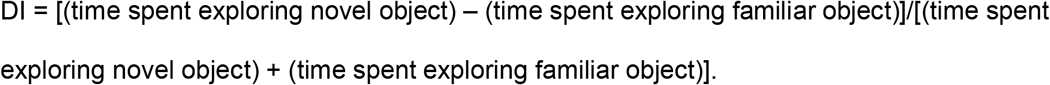

### Fiber photometry

A commercially available fiber photometry system, Neurophotometrics FP3002, was used. In brief, recording was accomplished by providing a 410 nm and 470 nm excitation light through the patch-cord for calcium-independent and calcium-dependent fluorescence emission from GCaMP7. Excitation power was adjusted to provide 75 μW of 410 nm and 470 nm light at the tip of the patch cord. Recordings were performed with the bonsai open-source software^52^ at 20 Hz. Analysis of the recorded traces was performed as previously described^53^. For recordings, mice were presented with either one inanimate object or one novel or one familiar mouse under a wire cup cage in an open arena. One item was present at a time and experimental mice were allowed to explore it for 5 min.

### Statistical analysis

We used Prism (Graphpad) for statistical analysis. Results presented in the figures are reported as the mean ± s.e.m. The statistical significance was tested by t-tests, paired t-tests or ANOVA (one-way, two-way or repeated measures) followed by Post-hoc Holm-Sidak’s multiple comparisons, as indicated. p<0.05 was considered significant.

## Author contributions

J.L-R. and S.A.S. conceptualized the research; J.L-R. carried out experiments and analysed data; C.A.S. carried out fiber photometry experiments. F.L. carried out rabies tracing experiments; J.L-R. wrote the original draft of the manuscript; J.L-R. and S.A.S. reviewed and edited the manuscript with input from all authors; S.A.S. and E.R.K. supervised the research and acquired funding.

## Acknowledgements

We thank members of the Siegelbaum laboratory for comments and discussions on the manuscript; L. M. Boyle for pilot calcium imaging experiments and H. S. Chen for helping with inmunostainings. This work was supported by R01-MH104602 and R01-MH106629 from N.I.H. to S.A.S.

## References

1. Kafkas, A. & Montaldi, D. How do memory systems detect and respond to novelty? Neurosci Lett 680, 60–68 (2018).

2. Strange, B. A., Witter, M. P., Lein, E. S. & Moser, E. I. Functional organization of the hippocampal longitudinal axis. Nat Rev Neurosci 15, 655–669 (2014).

3. Fernández, G. & Morris, R. G. M. Memory, Novelty and Prior Knowledge. Trends in Neurosciences 41, 654–659 (2018).

4. Hitti, F. L. & Siegelbaum, S. A. The hippocampal CA2 region is essential for social memory. Nature 508, 88–92 (2014).

5. Meira, T. et al. A hippocampal circuit linking dorsal CA2 to ventral CA1 critical for social memory dynamics. Nat Commun 9, 4163 (2018).

6. Oliva, A., Fernández-Ruiz, A., Leroy, F. & Siegelbaum, S. A. Hippocampal CA2 sharp-wave ripples reactivate and promote social memory. Nature 1–6 (2020) doi:10.1038/s41586-020-2758-y.

7. Donegan, M. L. et al. Coding of social novelty in the hippocampal CA2 region and its disruption and rescue in a 22q11.2 microdeletion mouse model. Nat Neurosci 1–11 (2020) doi:10.1038/s41593-020-00720-5.

8. Middleton, S. J. & McHugh, T. J. CA2: A Highly Connected Intrahippocampal Relay. Annual Review of Neuroscience 43, null (2020).

9. Alexander, G. M. et al. Social and novel contexts modify hippocampal CA2 representations of space. Nature Communications 7, 10300 (2016).

10. Watarai, A., Tao, K., Wang, M.-Y. & Okuyama, T. Distinct functions of ventral CA1 and dorsal CA2 in social memory. Current Opinion in Neurobiology 68, 29–35 (2021).

11. Chen, S. et al. A hypothalamic novelty signal modulates hippocampal memory. Nature 1–5 (2020) doi:10.1038/s41586-020-2771-1.

12. Eichenbaum, H., Yonelinas, A. P. & Ranganath, C. The Medial Temporal Lobe and Recognition Memory. Annu. Rev. Neurosci. 30, 123–152 (2007).

13. Reagh, Z. M. & Yassa, M. A. Object and spatial mnemonic interference differentially engage lateral and medial entorhinal cortex in humans. Proc Natl Acad Sci USA 111, E4264–E4273 (2014).

14. Connor, C. E. & Knierim, J. J. Integration of objects and space in perception and memory. Nature Neuroscience 20, 1493–1503 (2017).

15. Chevaleyre, V. & Siegelbaum, S. A. Strong CA2 Pyramidal Neuron Synapses Define a Powerful Disynaptic Cortico-Hippocampal Loop. Neuron 66, 560–572 (2010).

16. Sun, Q., Srinivas, K. V., Sotayo, A. & Siegelbaum, S. A. Dendritic Na+ spikes enable cortical input to drive action potential output from hippocampal CA2 pyramidal neurons. eLife https://elifesciences.org/articles/04551 (2014) doi:10.7554/eLife.04551.

17. Srinivas, K. V. et al. The Dendrites of CA2 and CA1 Pyramidal Neurons Differentially Regulate Information Flow in the Cortico-Hippocampal Circuit. J. Neurosci. 37, 3276–3293 (2017).

18. Kohara, K. et al. Cell type–specific genetic and optogenetic tools reveal hippocampal CA2 circuits. Nat Neurosci 17, 269–279 (2014).

19. McHugh, T. J. et al. Dentate gyrus NMDA receptors mediate rapid pattern separation in the hippocampal network. Science (New York, N.Y.) 317, 94–99 (2007).

20. Zhu, H. et al. Cre-dependent DREADD (Designer Receptors Exclusively Activated by Designer Drugs) mice. Genesis 54, 439–446 (2016).

21. Lisman, J. E. & Grace, A. A. The hippocampal-VTA loop: controlling the entry of information into long-term memory. Neuron 46, 703–713 (2005).

22. Dudman, J. T., Tsay, D. & Siegelbaum, S. A. A Role for Synaptic Inputs at Distal Dendrites: Instructive Signals for Hippocampal Long-Term Plasticity. Neuron 56, 866–879 (2007).

23. Basu, J. et al. Gating of hippocampal activity, plasticity, and memory by entorhinal cortex long-range inhibition. Science 351, aaa5694 (2016).

24. Kaifosh, P. & Losonczy, A. Mnemonic Functions for Nonlinear Dendritic Integration in Hippocampal Pyramidal Circuits. Neuron 90, 622–634 (2016).

25. Moser, E. I., Moser, M.-B. & McNaughton, B. L. Spatial representation in the hippocampal formation: a history. Nat Neurosci 20, 1448–1464 (2017).

26. Rowland, D. C., Roudi, Y., Moser, M.-B. & Moser, E. I. Ten Years of Grid Cells. Annual Review of Neuroscience 39, 19–40 (2016).

27. Tsao, A. et al. Integrating time from experience in the lateral entorhinal cortex. Nature 561, 57–62 (2018).

28. Montchal, M. E., Reagh, Z. M. & Yassa, M. A. Precise temporal memories are supported by the lateral entorhinal cortex in humans. Nature Neuroscience 22, 284–288 (2019).

29. Wang, C. et al. Egocentric coding of external items in the lateral entorhinal cortex. Science 362, 945–949 (2018).

30. Vandrey, B. et al. Fan Cells in Layer 2 of the Lateral Entorhinal Cortex Are Critical for Episodic-like Memory. Current Biology 30, 169–175.e5 (2020).

31. Li, Y. et al. A distinct entorhinal cortex to hippocampal CA1 direct circuit for olfactory associative learning. Nature Neuroscience 20, 559–570 (2017).

32. Bota, M., Sporns, O. & Swanson, L. W. Architecture of the cerebral cortical association connectome underlying cognition. Proc Natl Acad Sci USA 112, E2093–E2101 (2015).

33. Swanson, L. W. & Kohler, C. Anatomical evidence for direct projections from the entorhinal area to the entire cortical mantle in the rat. J. Neurosci. 6, 3010–3023 (1986).

34. Leung, C. et al. Activation of Entorhinal Cortical Projections to the Dentate Gyrus Underlies Social Memory Retrieval. Cell Reports 23, 2379–2391 (2018).

35. Sun, Q. et al. Proximodistal Heterogeneity of Hippocampal CA3 Pyramidal Neuron Intrinsic Properties, Connectivity, and Reactivation during Memory Recall. Neuron 95, 656–672.e3 (2017).

36. Chiang, M.-C., Huang, A. J. Y., Wintzer, M. E., Ohshima, T. & McHugh, T. J. A role for CA3 in social recognition memory. Behavioural Brain Research 354, 22–30 (2018).

37. Okuyama, T., Kitamura, T., Roy, D. S., Itohara, S. & Tonegawa, S. Ventral CA1 neurons store social memory. Science 353, 1536–1541 (2016).

38. Hasselmo, M. E. & Wyble, B. P. Free recall and recognition in a network model of the hippocampus: simulating effects of scopolamine on human memory function. Behav Brain Res 89, 1–34 (1997).

39. Robert, V. et al. Local circuit allowing hypothalamic control of hippocampal area CA2 activity and consequences for CA1. bioRxiv 2020.09.18.303693 (2020) doi:10.1101/2020.09.18.303693.

40. Karayiorgou, M., Simon, T. J. & Gogos, J. A. 22q11.2 microdeletions: linking DNA structural variation to brain dysfunction and schizophrenia. Nature Reviews Neuroscience 11, 402–416 (2010).

41. Piskorowski, R. A. et al. Age-Dependent Specific Changes in Area CA2 of the Hippocampus and Social Memory Deficit in a Mouse Model of the 22q11.2 Deletion Syndrome. Neuron 89, 163–176 (2016).

42. Benes, F. M., Kwok, E. W., Vincent, S. L. & Todtenkopf, M. S. A reduction of nonpyramidal cells in sector CA2 of schizophrenics and manic depressives. Biol Psychiatry 44, 88–97 (1998).

43. Knable, M. B. et al. Molecular abnormalities of the hippocampus in severe psychiatric illness: postmortem findings from the Stanley Neuropathology Consortium. Mol Psychiatry 9, 609–620, 544 (2004).

44. Gómez-Isla, T. et al. Profound loss of layer II entorhinal cortex neurons occurs in very mild Alzheimer’s disease. J Neurosci 16, 4491–4500 (1996).

45. Khan, U. A. et al. Molecular drivers and cortical spread of lateral entorhinal cortex dysfunction in preclinical Alzheimer’s disease. Nature Neuroscience 17, 304–311 (2014).

46. Kobro-Flatmoen, A., Nagelhus, A. & Witter, M. P. Reelin-immunoreactive neurons in entorhinal cortex layer II selectively express intracellular amyloid in early Alzheimer’s disease. Neurobiology of Disease 93, 172–183 (2016).

47. Leroy, F., Brann, D. H., Meira, T. & Siegelbaum, S. A. Input-Timing-Dependent Plasticity in the Hippocampal CA2 Region and Its Potential Role in Social Memory. Neuron 95, 1089–1102.e5 (2017).

48. Leroy, F. et al. A circuit from hippocampal CA2 to lateral septum disinhibits social aggression. Nature 564, 213–218 (2018).

49. Kogan, J. H., Franklandand, P. W. & Silva, A. J. Long-term memory underlying hippocampus-dependent social recognition in mice. Hippocampus 10, 47–56 (2000).

50. Yang, M. & Crawley, J. N. Simple Behavioral Assessment of Mouse Olfaction. Current Protocols in Neuroscience 48, 8.24.1–8.24.12 (2009).

51. Arbuckle, E. P., Smith, G. D., Gomez, M. C. & Lugo, J. N. Testing for Odor Discrimination and Habituation in Mice. JoVE (Journal of Visualized Experiments) e52615 (2015) doi:10.3791/52615.

52. Lopes, G. et al. Bonsai: an event-based framework for processing and controlling data streams. Front. Neuroinform. 9, (2015).

53. Martianova, E., Aronson, S. & Proulx, C. D. Multi-Fiber Photometry to Record Neural Activity in Freely-Moving Animals. JoVE (Journal of Visualized Experiments) e60278 (2019) doi:10.3791/60278.

