## Extended Data Fig. 1 for "A direct lateral entorhinal cortex to hippocampal CA2 circuit conveys social information required for social memory"

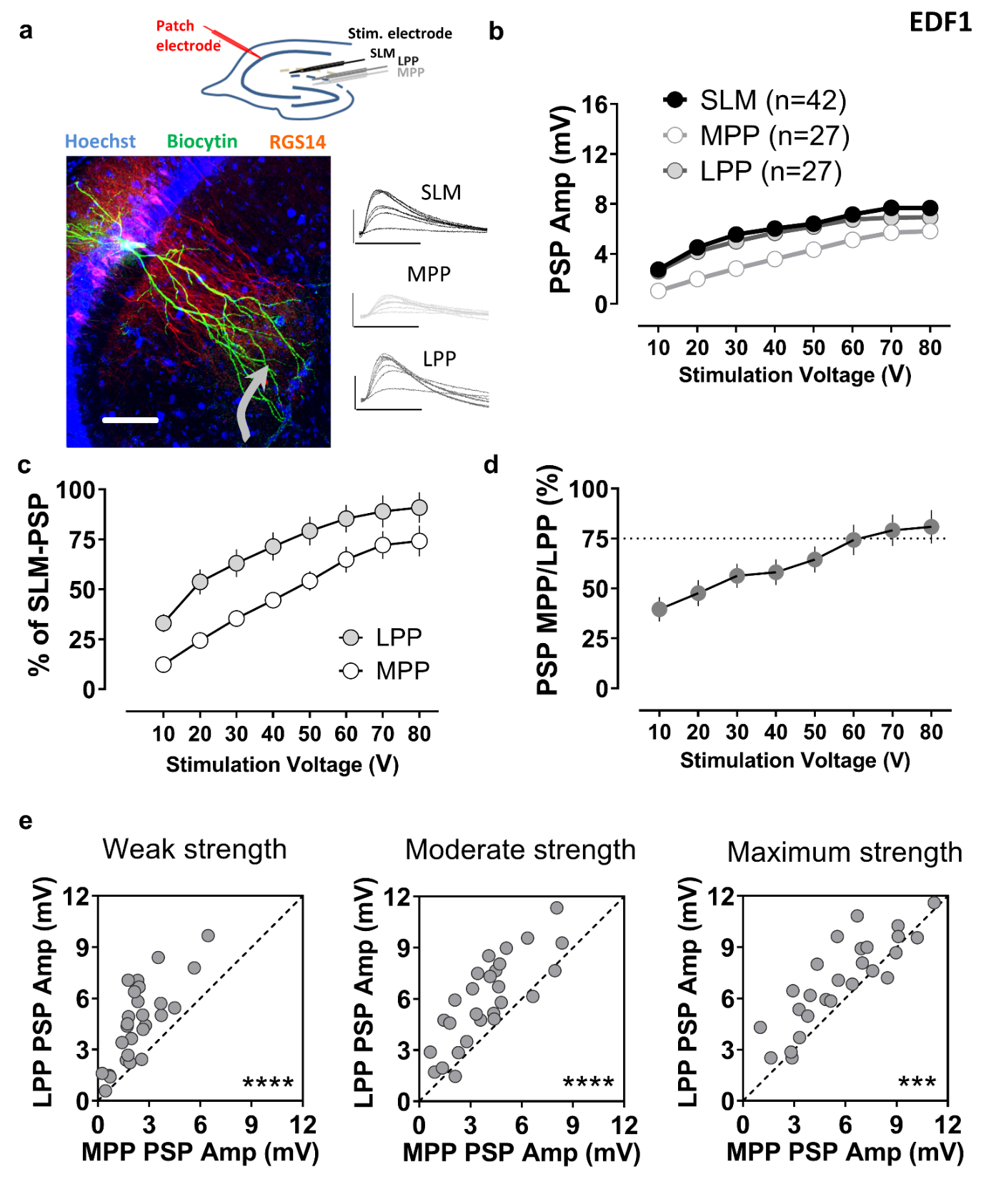
**Extended Data Figure 1. Electrical stimulation of the lateral perforant path evokes a larger intracellular synaptic depolarization of dorsal CA2 pyramidal neurons compared to stimulation of medial perforant path.** **a,** Schematic showing the placement of a stimulating electrode (black) in the stratum lacunosum moleculare (SLM) to globally activte entorhinal axons or in the outer (LPP) or middle (MPP) molecular layer of the dentate gyrus to selectively stimulate lateral or medial perforant path (LPP or MPP). Synaptic responses were measured using patch clamp recordings from CA2 pyramidal neurons. An image of a patch-clamped biocyting-filled CA2 pyramidal cell and the corresponding voltage responses to stimulation in the indicated regions. **b,** Electrical stimulation at any location evoked a large postsynaptic potential in CA2 pyramidal neurons. The LPP-evoked response was larger than the MPP-evoked response, expressed as percentage of the SLM-evoked response (**c**), as a ration of the MPP/LPP response (**d**) or as voltage amplitudes (**e**) over a range of stimulation strengths. Scale bars: **a**: 100 µm and 5 mV/25 ms. ****: p<0.0001, ***: p<0.0001 paired t-test.
