## Extended Data Fig. 2 for "A direct lateral entorhinal cortex to hippocampal CA2 circuit conveys social information required for social memory"

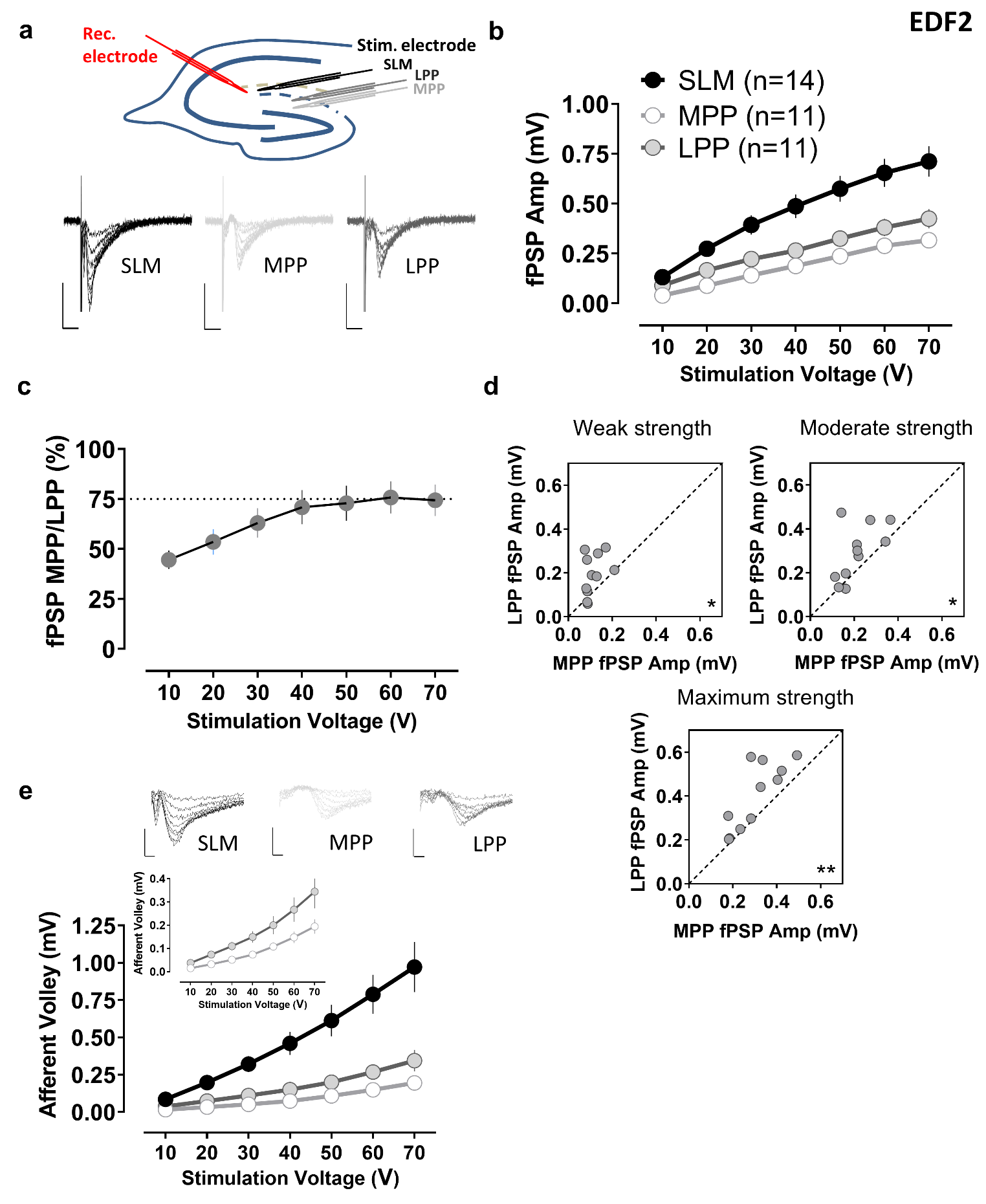

**Extended Data Figure 2. Electrical stimulation of the lateral perforant path evokes a larger extracellular field potential in SLM of dorsal CA2 region compared to stimulation of medial perforant path .** **a,** Schematic showing the placement of stimulating electrodes as in Extended Data Figure 1, while extracellularly recording from CA2 distal dendrites in SLM. The insets show representative field potentials. **b,** Electrical focal stimulation in SLM evoked a large field potential in CA2 distal dendrites. The LPP-evoked response was significantly larger than the MPP-evoked response at all stimulation strengths (**c, d**). **e,** The extracellular presynaptic fiber volley in response to a stimulating pulse, which reflects the number of activated axons, was also larger in LPP than in MPP, suggesting a larger number of synaptic contacts. Scale bars: **a**: 0.5 mV/5 ms, **e**: 0.5 mV/1 ms. *: p<0.05, **: p<0.01 paired t-test.
