## Extended Data Fig. 3 for "A direct lateral entorhinal cortex to hippocampal CA2 circuit conveys social information required for social memory"

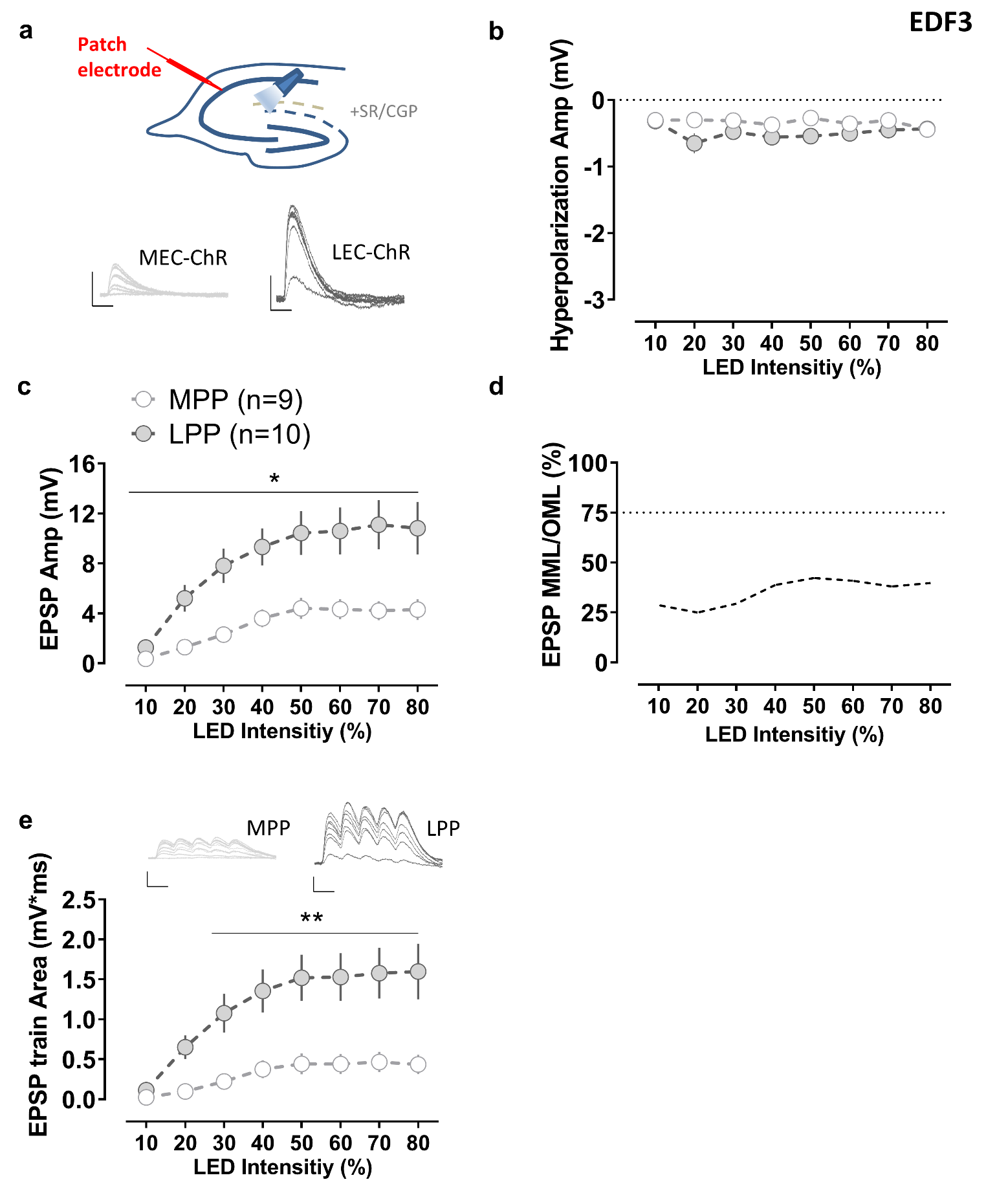

**Extended Data Figure 3. Optogenetic activation of lateral entorhinal cortex evokes larger excitatory postsynaptic potentials in CA2 pyramidal cells compared to activation of medial entorhinal cortex.** **a,** An AAV was injected to express ChR2 in the medial (MEC) or the lateral entorhinal cortex (LEC). All recordings were done in the presence of GABA receptor blockers to isolate the pure excitatory response (**b**). Pulses of blue light were shone on the stratum lacunosum moleculare while intracellularly recording from CA2 pyramidal neurons. **c, d,** Photostimulation of ChR2-expressing terminals in the stratum lacunosum moleculare evoked a large excitatory postsynaptic potential in CA2 pyramidal neurons for both, MEC and LEC groups. The LEC-evoked response was significantly larger than the MPP-evoked one over a range of stimulation strengths for a single light pulse and for a short train of optical stimuli (**e**). Scale bars: **a, e**: 5 mV/25 ms. **: p<0.01 Holm-Sidak's post hoc test after two-way mixed-design ANOVA.
