## Extended Data Fig. 4 for "A direct lateral entorhinal cortex to hippocampal CA2 circuit conveys social information required for social memory"

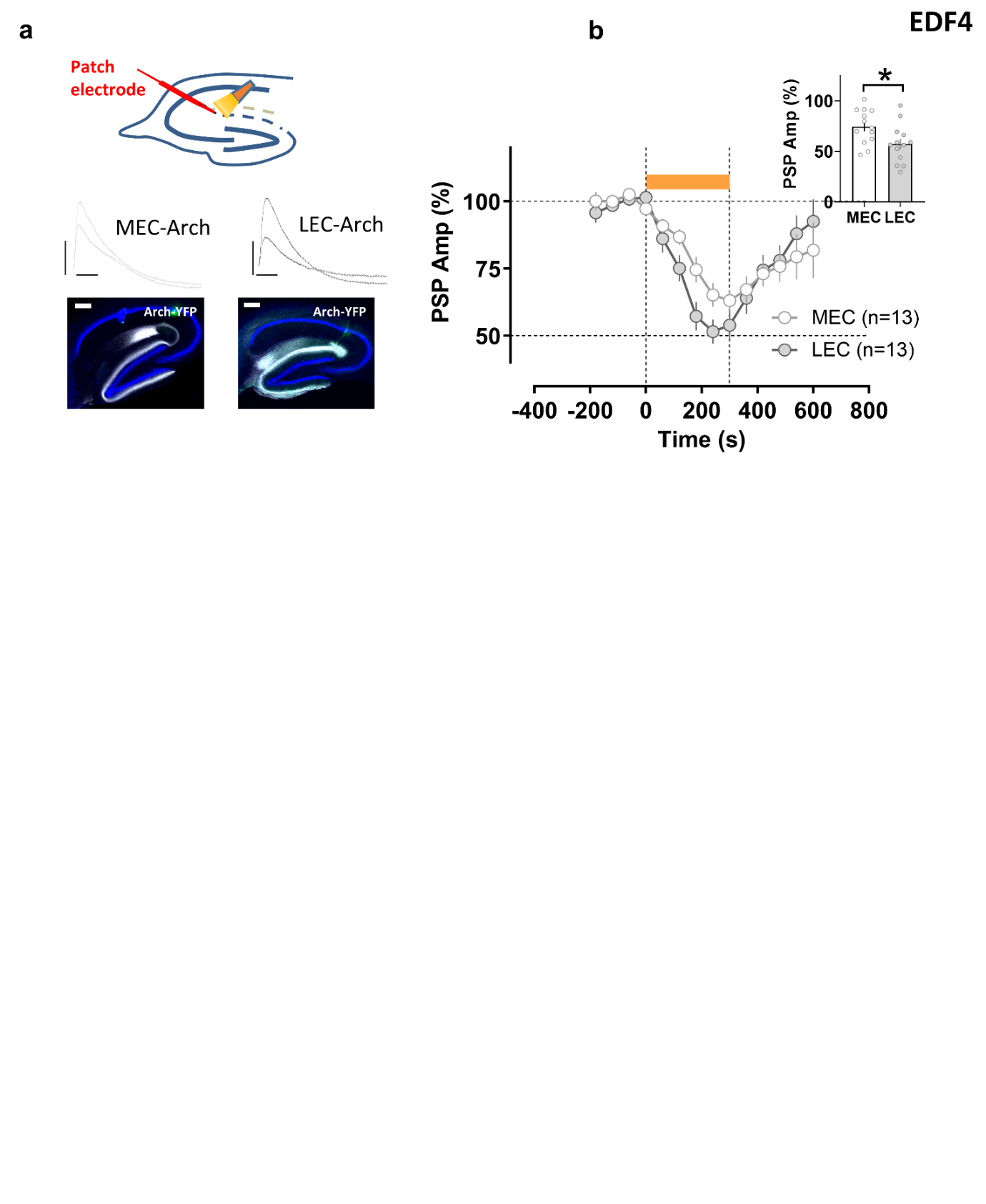

**Extended Data Figure 4. Optogenetic inhibition of inputs to CA2 reveals larger contribution of the lateral compared to medial entorhinal cortex. a,** An AAV was injected in medial (MEC) or lateral entorhinal cortex (LEC) to express Arch. Illumination of the stratum lacunosum moleculare (SLM) with yellow light was used to assess the effect of optogenetic inhibition of LEC or MEC inputs on the postsynaptic depolarization in CA2 pyramidal cells evoked by electrical stimulation using an electrode in SLM. **b,** Temporal course of evoked responses. Yellow light was on for 300 s as shown in the graph. The inset shows recorded responses after 180 s of continuos illumination. Scale bars: 5 mV/25 ms and 200 µm. *: p<0.05 Holm-Sidak's post hoc test after two-way mixed-design ANOVA.
