## Extended Data Fig. 5 for "A direct lateral entorhinal cortex to hippocampal CA2 circuit conveys social information required for social memory"

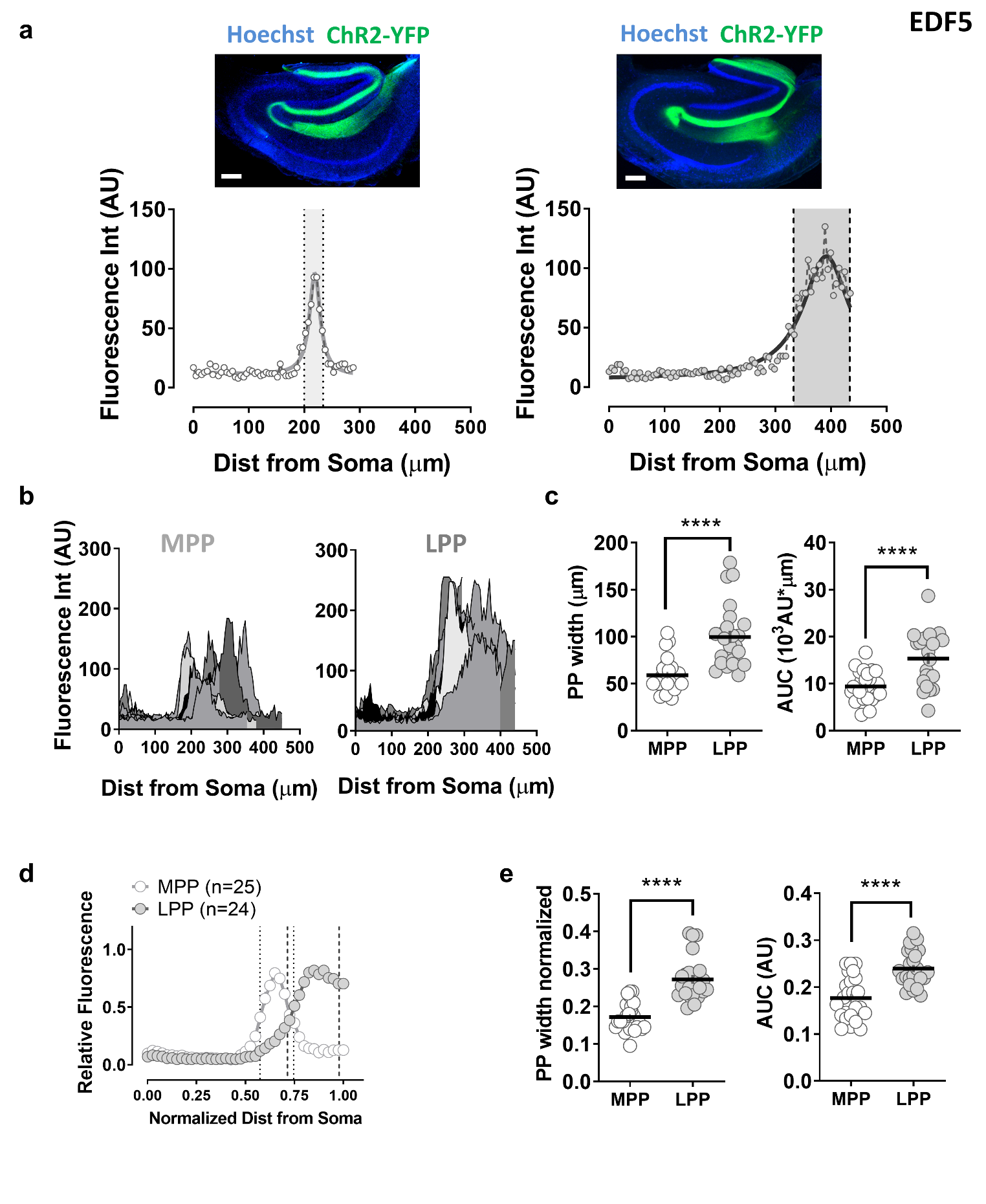

**Extended Data Figure 5. Lateral entorhinal cortex axons (LPP) occupy a larger area than medial entorhinal cortex ones (MPP) in the stratum lacunosum moleculare (SLM) of CA2.** **a,** Example images of medial, left, or lateral, right, entorhinal axons in hippocampus labelled by injecting AAV to express ChR2-YFP in MEC or LEC, respectively. Traces below the images show the intensity profiles of entorhinal axon fluorescence for ChR2-YFP along the CA2 radial axis, from the pyramidal cell layer up to SLM. **b,** Intensity profiles of all analysed slices. **c,** LPP occupies a larger area in CA2 SLM than MPP. **d,** Intensity profiles with normalized values of fluorescence and distance. **e**, LPP has a significantly larger width in CA2 SLM than MPP. Scale bar: **a**: 200 µm. ****: p<0.0001 t-test.
