## Extended Data Fig. 6 for "A direct lateral entorhinal cortex to hippocampal CA2 circuit conveys social information required for social memory"

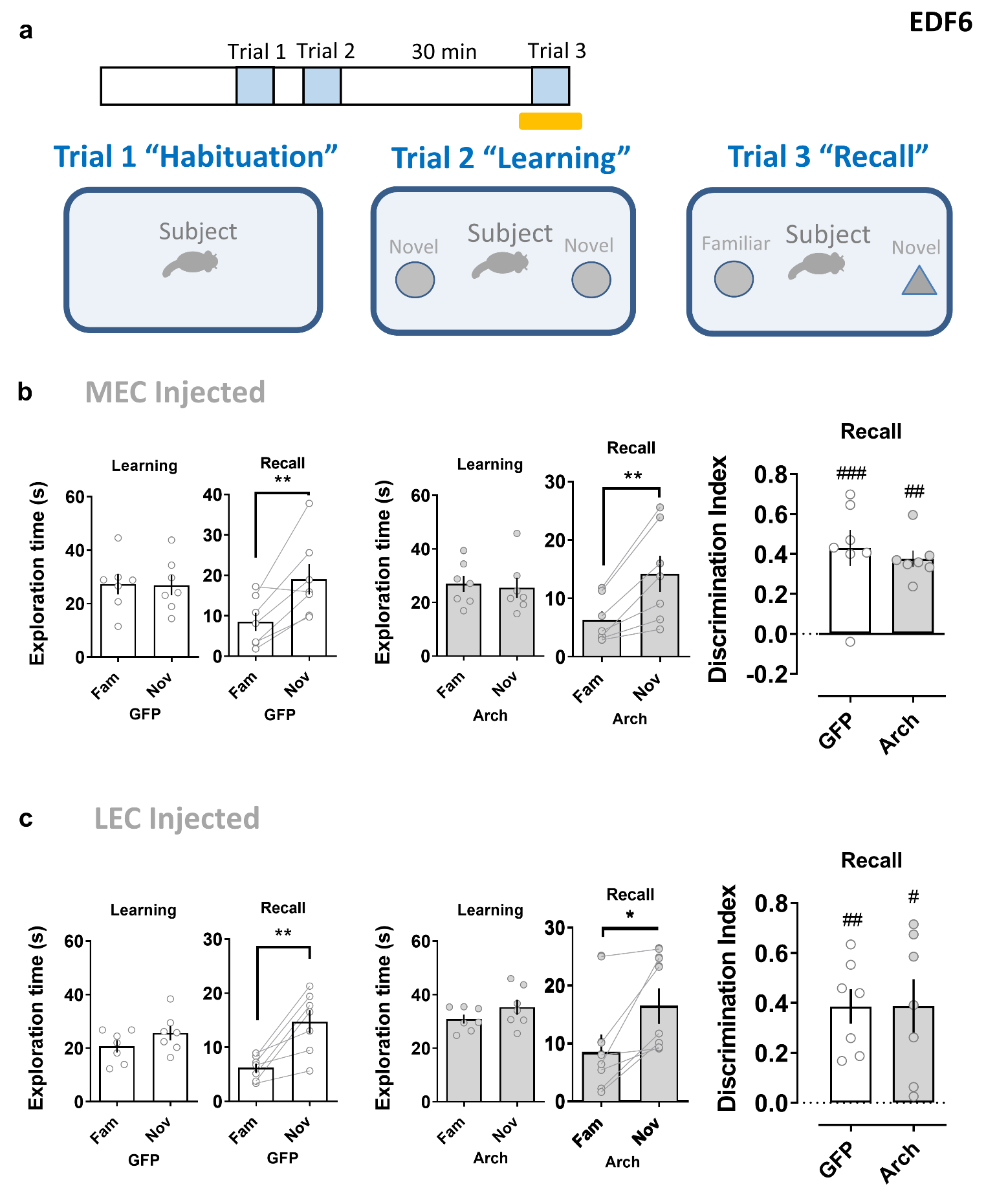

**Extended Data Figure 6. Disrupting the entorhinal input to dorsal CA2 does not alter novel object recognition. a**, Schema of the novel object recognition task, which is analogous to the two-chice social memory test. Shining yellow light on entorhinal cortex inputs in dorsal CA2 during the recall phase (trial 3) did not significantly alter the performance of animals previously injected with an Arch-expressing AAV in MEC (**b**) or LEC (**c**), compared to control groups injected with GFP-expressing AAV. #: p<0.05, ##: p<0.01, ###: p<0.001 one-sample t-test against “0”. *: p<0.05, **: p<0.01 paired t-test.
