## Extended Data Fig. 7 for "A direct lateral entorhinal cortex to hippocampal CA2 circuit conveys social information required for social memory"

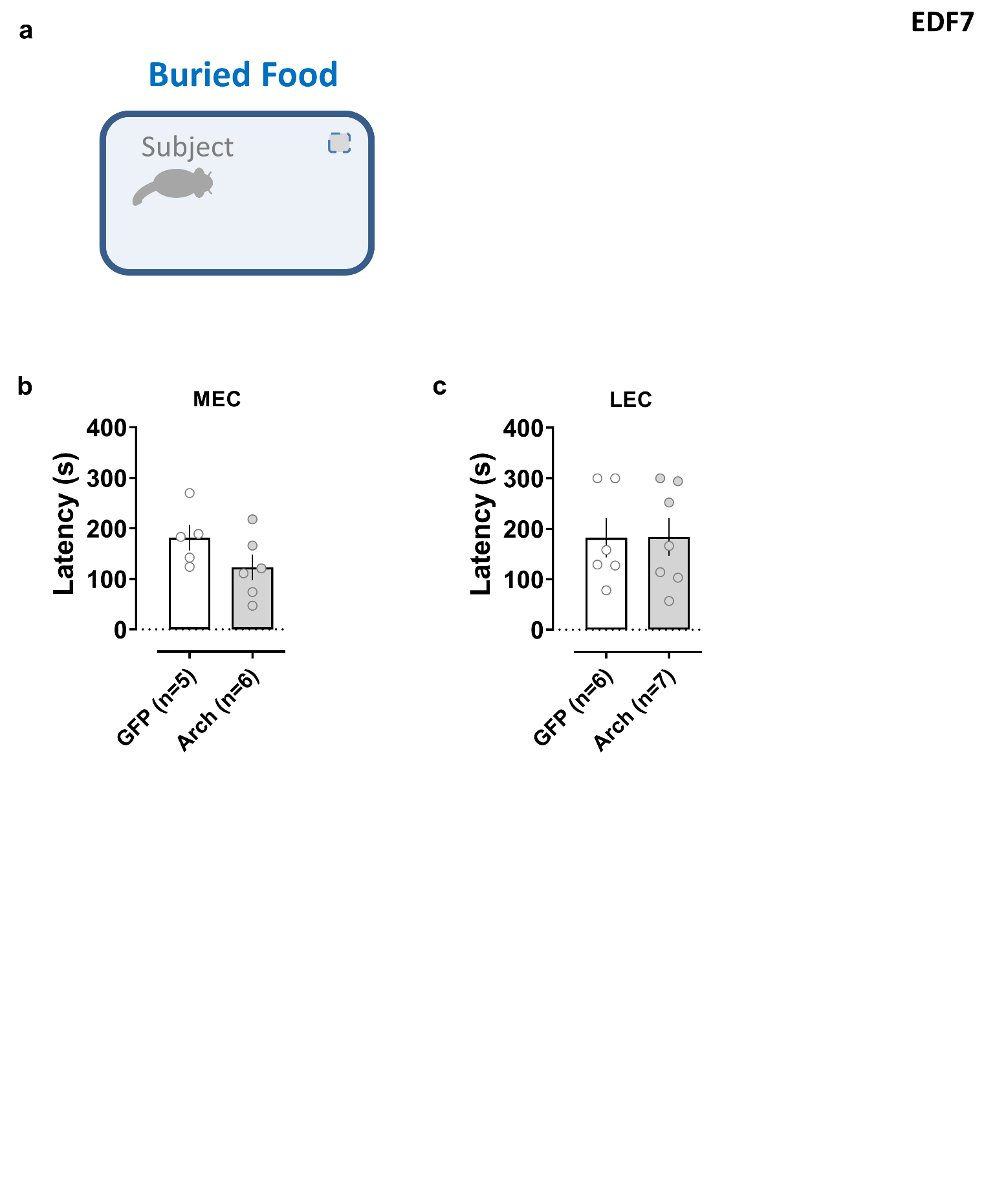

**Extended Data Figure 7. Disrupting the entorhinal input to dorsal CA2 does not alter olfactory task performance.** **a**, Food deprived animals searched to find a buried pellet of food. Shining yellow light on entorhinal cortex inputs in dorsal CA2 did not significantly change the performance of animals previously injected with an Arch-expressing AAV in MEC (**b**) or LEC (**c**), compared to GFP-expressing control groups.
