## Extended Data Fig. 8 for "A direct lateral entorhinal cortex to hippocampal CA2 circuit conveys social information required for social memory"

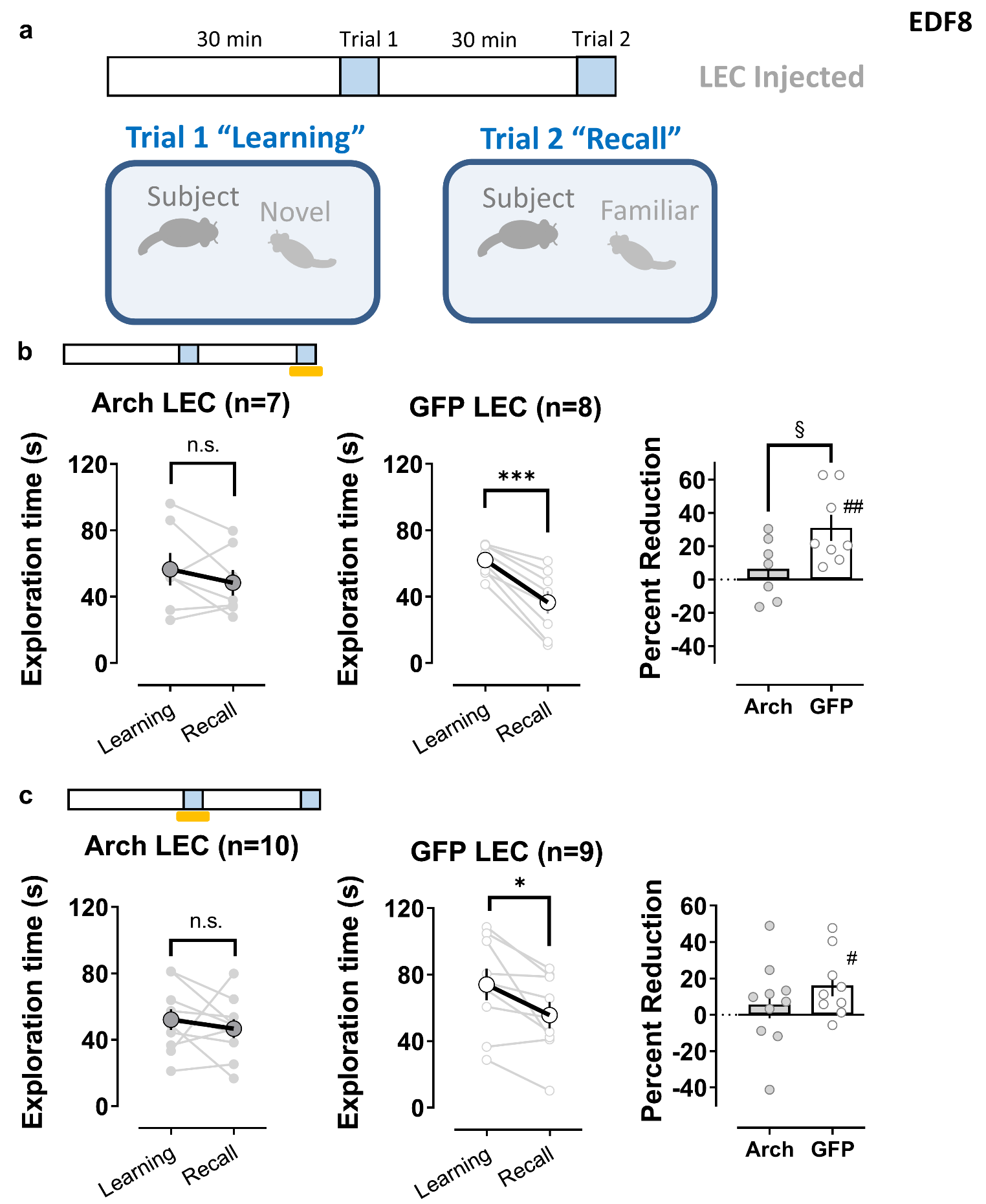

**Extended Data Figure 8. Disrupting the lateral entorhinal input to dorsal CA2 impairs social memory in the direct interaction task. a**, Schema of the direct interaction social memory task. A male adult subject mouse was placed in a clean cage for a 30min habituation period. A novel juvenile male mouse was then introduced in the cage and the subject mouse was allowed to explore it for 2 min (trial 1, learning). The juvenile was removed and after a 30 min interval, the same mouse is reintroduced in trial 2. Social memory is manifest as a decrease in exploration of the now familiar juvenile in trial 2 compared to trial 1. Shining yellow light on lateral entorhinal cortex (LEC) inputs in dorsal CA2 either during the recall phase (trial 2) (**b**) or during the learning phase (trial 1) (**c**), impairs the performance of animals previously injected with an Arch-expressing AAV in LEC compared to the control group expressing GFP in LEC. §: p<0.05 t-test. #: p<0.05, ###: p<0.001 one-sample t-test against “0”. *: p<0.05,***: p<0.001, paired t-test.
