## Extended Data Fig. 9 for "A direct lateral entorhinal cortex to hippocampal CA2 circuit conveys social information required for social memory"

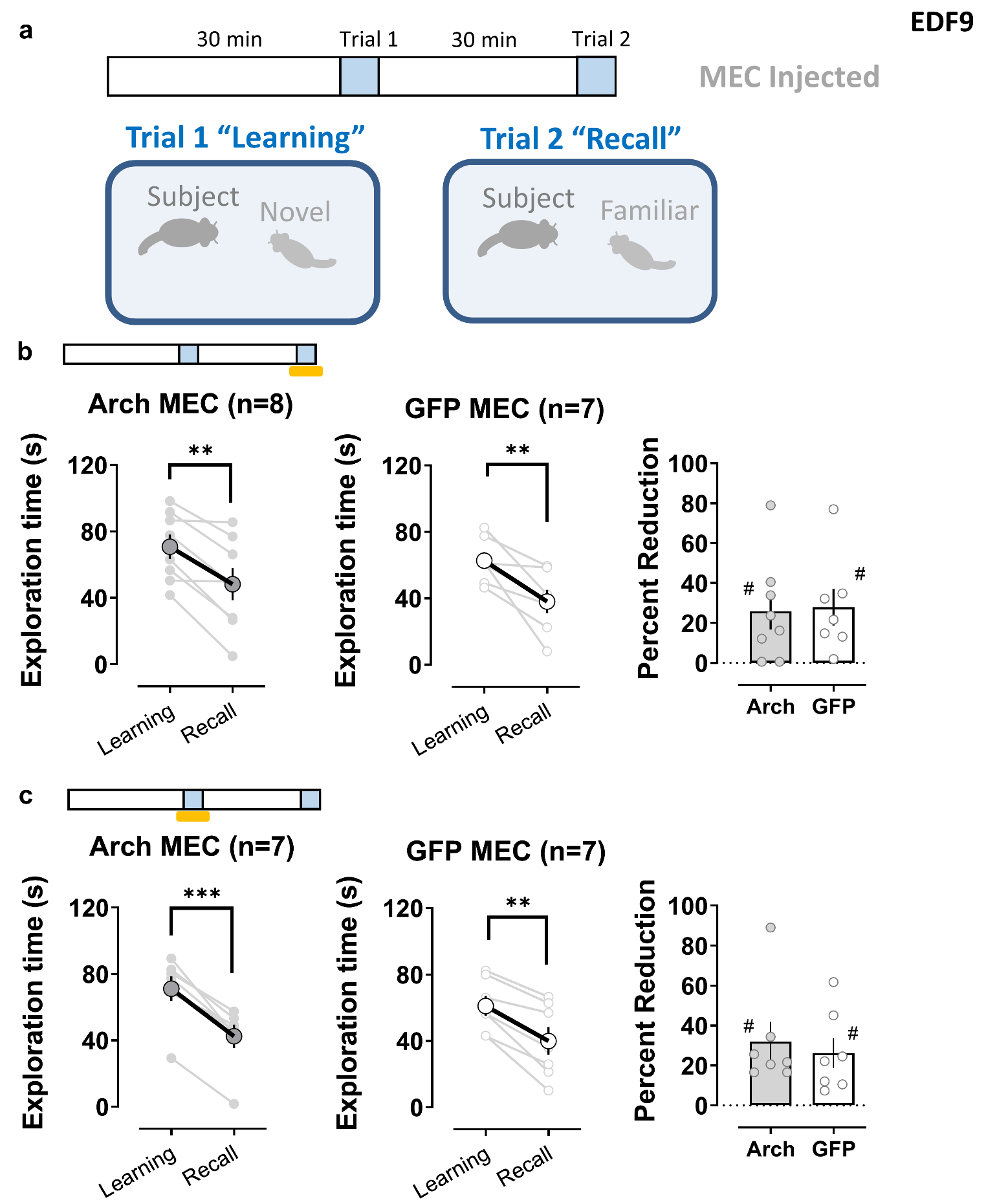

**Extended Data Figure 9. Disrupting the medial entorhinal input to dorsal CA2 does not influence performance in the direct interaction social memory task. a**, Schema of the direct interaction task consisting of two trials. Shining yellow light on medial entorhinal cortex (MEC) inputs in dorsal CA2 either during the recall phase of the task (trial 2) (**b**) or during the learning phase (trial 1) (**c**), does not significantly change the performance of animals previously injected with an Arch-expressing AAV in MEC compared to a control GFP-expressing group. ##: p<0.01, ###: p<0.001 one-sample t-test against “0”. **: p<0.01,***: p<0.001, paired t-test.
