## Extended Data Fig. 10 for "A direct lateral entorhinal cortex to hippocampal CA2 circuit conveys social information required for social memory"

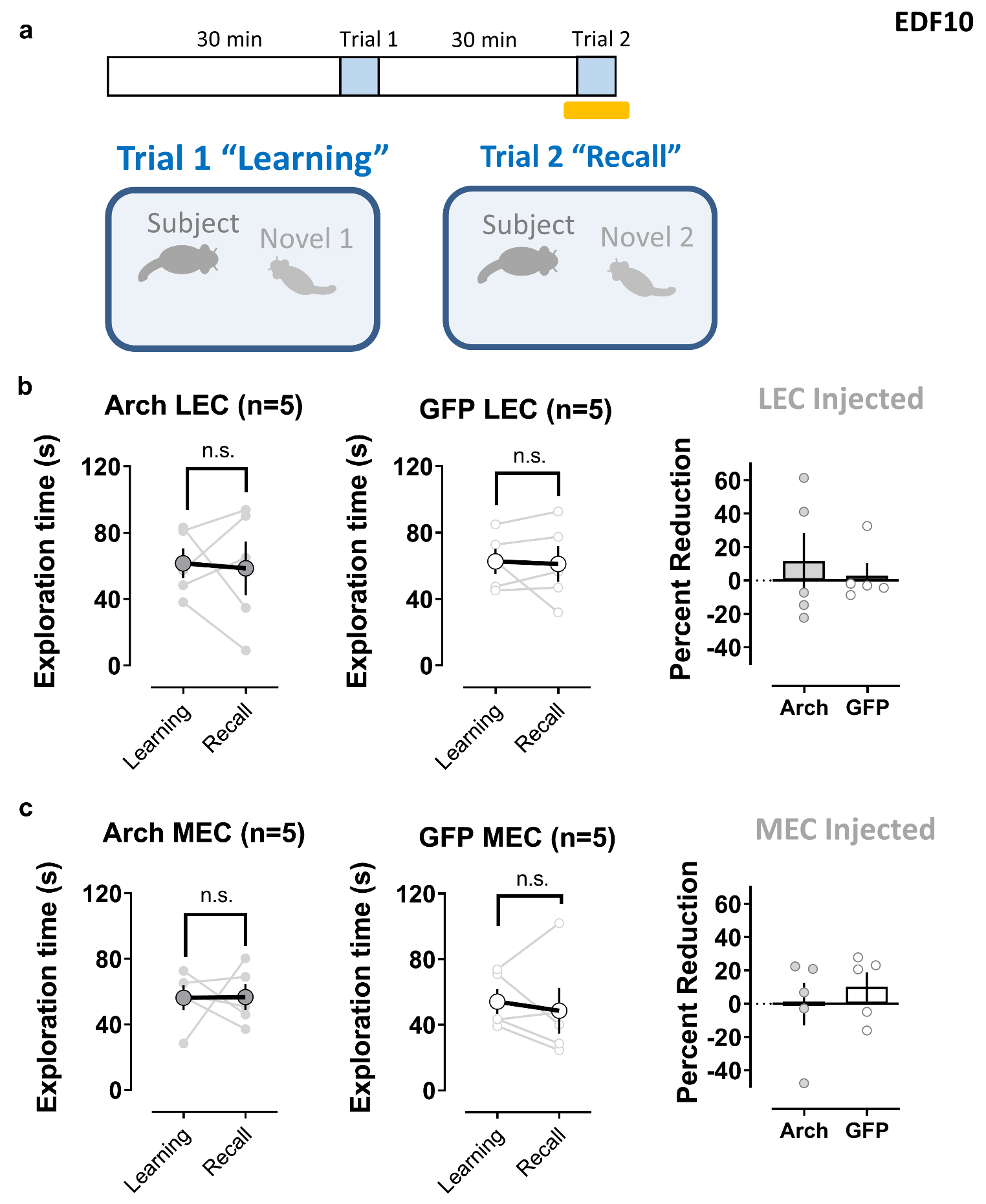

**Extended Data Figure 10. Decreased exploration time during the recall trial in the direct interaction task does no result from fatigue or lack of motivation for social exploration of the subject mice. a**, Schema of the behavioral task, a variant of the direct interaction task, consisting of two trials in which a second novel juvenile male was introduced in trial 2. Mice normally show equal exploration of the two novel juveniles, indicating that the decrease in exploration when the same juvenile encountered in trial 1 is reintroduced in trial 2 reflects social memory of the now familiar mouse, rather than fatigue or lack of motivation to explore in the second trial. Shining yellow light on entorhinal cortex inputs in dorsal CA2 during the recall phase (trial 2) did not significantly change the performance of animals previously injected with an Arch-expressing AAV in LEC (**b**) or MEC (**c**), compared to GFP-expressing control groups.
